# Translation fidelity mechanisms and their changes in aging

**DOI:** 10.64898/2026.09.24.754133

**Authors:** Mauricio Aguilar Rangel, Casey Powers, Jaeyoon Lee, Haley E. Tarbox, Neil B. Wood, Stephen D. Fried, Stephen Quake, Joseph D. Puglisi

## Abstract

Protein errors (amino acid misincorporations) are associated to protein misfolding and misfunction, and lower levels correlate with longer lifespan ^1^. Despite their prevalence, proteome-wide tracking of misincorporations in eukaryotes is challenging, with most insights being recent and coming from the mining of extensive proteomics data sets ^2–5^. These experimental challenges have precluded the direct *in vivo* probing and full understanding of the bases of fidelity in higher eukaryotes, and their influence in complex processes such as aging. Here we measured proteome-wide misincorporation frequencies in two experimentally tractable eukaryotic systems, yeast and mouse, using mass spectrometry. We find translation fidelity is organism and organ specific. The measured error frequencies correlate with codon:anticodon pairing thermodynamics and tRNA pool composition, which can have synergistic or opposing effects on fidelity depending on organism or tissue type. Most identified protein errors carry a negative fitness burden. Aging experiments in our two systems reveal that, with age, error frequencies increase in post-mitotic cells, but not so in mitotic ones. These changes correlate with remodeling of the tRNA pool, which we identify as a crucial regulator of translation fidelity during lifespan.

## Introduction

Proteins compose essential machines, enzymes and architecture of the cell. Thus, there is a constant need for the synthesis of new proteins. This demand is met by the ribosome-mediated translation of messenger RNAs (mRNAs) into proteins (herein translation). To ensure protein functionality, translation requires high fidelity. However, this process has the highest error rate of all gene expression steps, with an estimated 1 in 10,000 amino acid misincorporation frequency ^2,3,6–10^, with bacteria having higher error frequencies than eukaryotes ^2,9,10^. This error frequency has been proposed to result from a necessary trade-off between translation speed and fidelity, because the ribosome must translate fast enough to populate the cellular proteome while doing so accurately enough to preserve function ^11–16^. Protein demand is organism-dependent and therefore there is not a unique, universal translation error rate ^2,9,10,17^. Translation fidelity is tunable; it varies among organisms and can also be altered with drugs or ribosomal mutations ^18–23^. Intriguingly, variation in translation fidelity, whether natural or induced, strongly correlates with lifespan and aging, where higher fidelity is associated with longer lifespan ^1,24,25^.

The 64 codon (3 nucleotide) molecular alphabet for translation is substantially larger than the 4 nucleotide counterparts used in replication or transcription ^26–28^. Concomitantly, the number of ribosomal substrates, transfer RNAs (tRNAs), is large and diverse ^29^, creating a rich landscape of potential error mechanisms. tRNAs interact with mRNA codons in the ribosomal decoding center through their anticodon stem-loops (ASLs) and, during the initial stages of decoding, a codon can be sampled by any available tRNA as a ternary complex (TC) with EF-Tu/EF1A and GTP based solely on its concentration in the pool ^30,31^. The ribosome has evolved a complex multi-step mechanism to distinguish and favor correct codon-anticodon interactions ^32,33^, referred to as cognate pairs. Cognate tRNAs deliver the correct amino acid as a ternary complex (TC) by pairing to the mRNA with an energetically favorable mini-helix geometry facilitated by strictly-enforced Watson-Crick basepairing interactions, however wobble pairing at the 3^rd^ codon position is allowed ^34,35^. In contrast, interactions with unfavorable free energies result in codon-anticodon distorted geometries that cannot be easily accommodated within the decoding center ^34,35^, leading to shorter residency times, fast initial rejection, as well as subsequent rejection through kinetic proofreading ^36–38^. The anticodons of tRNAs that deliver the wrong amino acid, are classified into near- and non-cognate, with the former having one mis-paired nucleotide and the latter two or more ^31^. Discrimination of cognate and near/non-cognate tRNAs for translation fidelity thus kinetically magnifies the energetic differences of cognate and near-cognate codon-anticodon interactions ^38–40^.

The cellular tRNA pool can be as complex as hundreds of different isodecoders at different concentrations ^41^, all capable of sampling any codon awaiting decoding by the ribosome as a TC. Therefore, decoding is a competition between cognate- and near- and non-cognate tRNAs ^42^. Previous studies using reporter systems have shown how this competition regulates amino acid misincorporation for a few tRNAs and codons ^9,10^. It has also been shown that the tRNA pool can rapidly change in response to different cellular stresses ^43^, making the cognate vs. near-cognate tRNA competition malleable. Thus, understanding translation fidelity *in vivo* requires simultaneous monitoring of the cellular tRNA pool and translation errors across the entire proteome. Here, we employ state of the art mass spectrometry proteomics to track, proteome-wide, translation error rates in two eukaryotic systems, yeast and mouse. We establish how the tRNA pool regulates translation fidelity and then use structural and machine learning analyses to understand the functional implications of translation errors in the proteome. Finally, we show, in yeast and mouse aging models, how aging induces an imbalance in the cellular levels of cognate and near-cognate tRNAs, resulting in altered translation error frequencies.

## Results

### Trapped ion mobility mass spectrometry enables proteome-wide fidelity estimation in eukaryotic proteomes

We sought to measure translation fidelity across eukaryotic proteomes. Traditional reporter systems rely on fluorescent or luminescent readouts of amino acid misincorporation at catalytic residues of an exogenous protein ^1,9,10^. These systems, despite their importance, report on a few codons and tRNAs only, in the context of an exogenous mRNA. However, translation is context-dependent, with the ribosome displaying an uneven speed of translation correlated to tRNA abundance, codon usage, and amino acid properties ^44^. Hence, ideally fidelity should be evaluated on endogenous mRNAs in a proteome-wide manner. Elegant foundational work by Mordret and coworkers applied LC-MS/MS proteomics to measure error frequencies across the simple *Escherichia coli* proteome ^2^, however similar approaches in eukaryotic proteomes have been limited by the depth of coverage needed and the inherent bias in mass spectrometry favoring abundant proteins. Thus, depth and sensitivity must be increased to capture rare amino acid misincorporation events, which are inherently rare.

To solve these challenges, we exploited the capabilities of the new generation of timsTOF instruments, in which parallel accumulation of ions allows for a near-perfect duty cycle, while serial fragmentation enables a sequencing speed nearing 300Hz. We reasoned that these features should improve the detection of rare misincorporation events. We digested the total cellular extract of *S. cerevisiae* (which has a small proteome for a eukaryote, ∼6000 proteins) with trypsin, performed high-pH off-line fractionation, and analyzed the fractions by LC-MS/MS on a timsTOF Ultra (Fig. 1B). We ensured no sequencing bias in the data, from oversampling of abundant proteins, by comparing the codon composition of the identified peptides against the expected codon composition of the translatome based on ribosome profiling data (R = 0.96, Fig. S1A). Spectra were sequenced with PEAKS Online and searched against the yeast proteome to annotate wild-type (WT) spectra. Candidate amino acid misincorporations were identified with SPIDER ^45^, followed by final identification and filtering with our custom pipeline, available as an R package (Table S1, see *Data availability*). The identified WT and misincorporation-harboring spectra were mapped back to their corresponding codon positions, and we calculated codon error frequencies as the quotient of misicorporation over WT codon counts. We were able to measure 39 codon misincorporation frequencies (Fig. 1C) and, given the depth of the data, could also measure stop codon readthrough (SCR) frequency (Fig. 1D). SCR frequencies follow the permissiveness UGA>UAG>UAA (Fig. S1B), in agreement with yeast measurements based on ribosome profiling data ^46,47^. Technical validation of our pipeline was twofold: first, when using the *S. cerevisiae s288c* proteome as reference in our analysis, which harbors 37 amino acid mutations respect to the actual measured strain (BY4743, Table S2), several of these genomic differences between strains were successfully identified as putative amino acid substitutions (Table S1). Second, a different spectral assignment approach, utilizing Open Search from MSFragger ^48^ as proposed by Landerer et al. for Orbitrap data ^3^, resulted in a well-correlated set of error estimates (Fig. S1C). To validate the biological origin of the mapped mutations, we performed the same experiment with yeast grown in the presence of paromomycin, which stabilizes near-cognate pairings at the ribosome decoding center, thus increasing the probability of near-cognate misincorporations and a higher SCR frequency ^46,49^. Paromomycin increased total error frequency by 47%, driven by an increase in near-cognate error at most measured codons (Fig. S1D-G); it also increased SCR frequency for all stop codons, causing an overall 58% increase in SCR (Fig. S1H,I). These results confirm that the substitutions detected by this method are related to ribosomal misincorporation, and not due to genetic variants or errors during transcription (which are thought to occur at considerably lower frequencies). We next tested our approach with a more complex mammalian proteome, mouse, which has ∼20,000 proteins. Using total extract from mouse liver we were able to quantify 35 codon error frequencies (Fig. 1C, Table S1) and the corresponding SCR error (Fig. 1D,S1J). Remarkably, average per codon misincorporation frequency in mouse is lower and with a tighter distribution (2e-05±2.1e-05) than in yeast (3.2e-05±4.4e-05).

**Figure 1.**
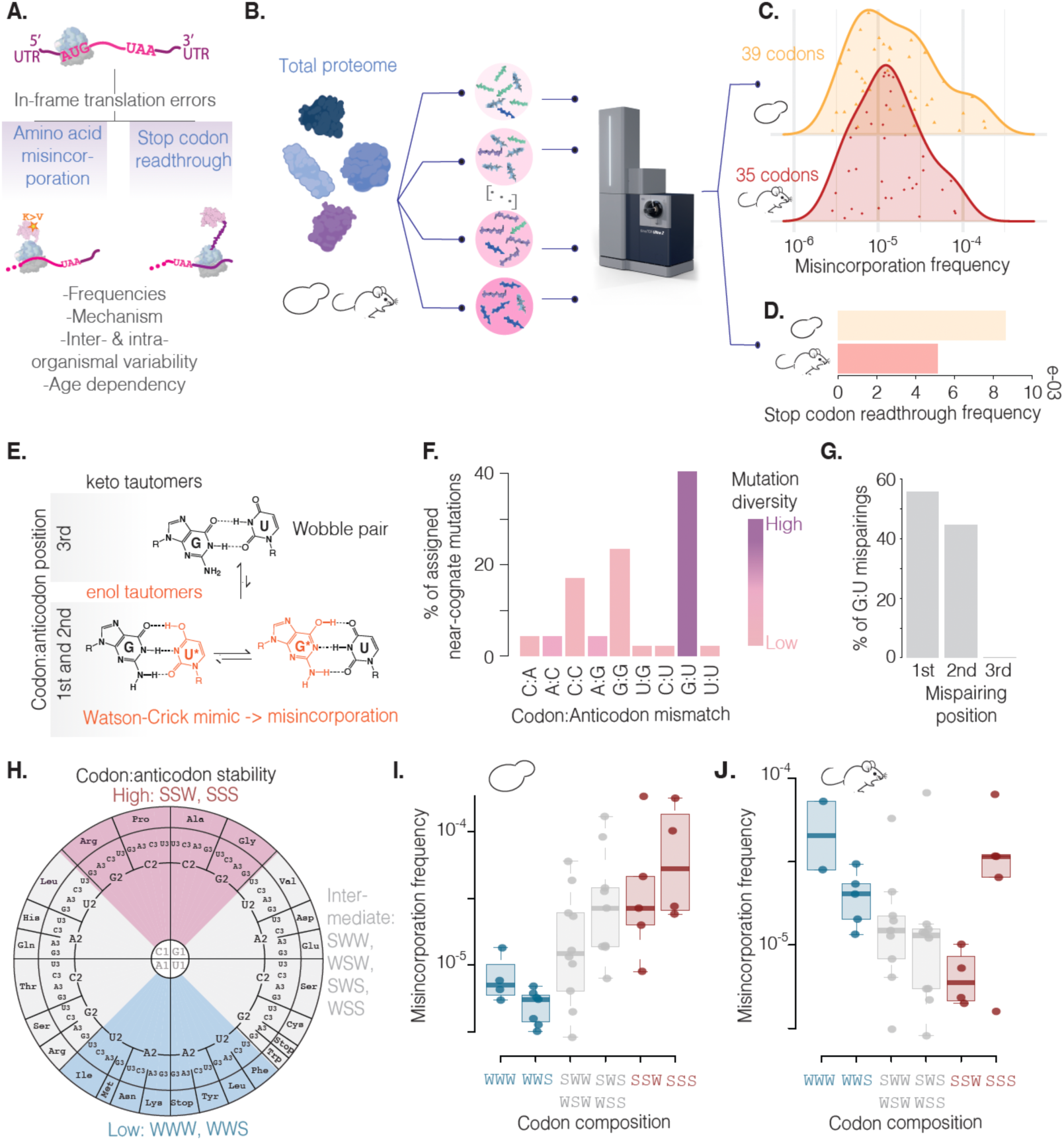
Translation error mapping in eukaryotic proteomes and their molecular signatures. A) Types of protein translation errors happening in-frame. B) Schematic of the utilized sample processing for protein error identification. C) Distributions of the measured codon misincorporation frequencies in yeast and mouse. D) Measured total stop codon readthrough frequencies in yeast and mouse. E) Schematic depicting the equilibria between G:U wobble pairing and the corresponding keto-enol states. F) Types and percentages of the different codon:anticodon mispairings assigned to the mapped yeast amino acid misincorporations. G) Percentages of misincorporation-assigned G:U mispairings at each codon:anticodon position, corresponding to yeast misincorporations. H) The energy-based organization of the genetic code proposed by Grosjean and Westhof, with IUPAC nomenclature for the different energy-based codon categories. I-J) Measured codon misincorporation frequencies binned according to the corresponding energy-based category of the codon, for yeast (I) and mouse (J).

### Translation fidelity correlates with codon-anticodon stability

Codon translation error frequencies span a wide range of values in both analyzed organisms, from 10^-6^ to 10^-4^, and thus we set out to characterize what distinguishes codons with lower or higher error frequencies. We determined the codon-anticodon mispairing position for the observed mutations in both organisms, and surprisingly most occur at the first codon-anticodon position, with frequencies close to zero for the third one (Fig. S2A,B). When the mispairing could be unambiguously assigned to a single near-cognate tRNA, G:U mispairings represent the most frequent misincorporation source, with G:G in second place, for both mouse and yeast (Fig. 1F, S2C). G:U pairing is generally permitted at the 3^rd^ position as a wobble pair, but not at the 1^st^ due to steric clash from wobble pairing geometry. Alternatively, nucleic acid bases can adopt two different *keto-enol* tautomeric forms and, despite the *keto* form being overly predominant, the *enol* tautomer (N*) allows G*:U and G:U* to adopt a stable, 3-hydrogen-bonded, G:C-like geometry ^50^ (Fig. 1E). Codon-anticodon pairs containing those tautomers structurally resemble cognate interactions ^51,52^. Moreover, enol tautomers comprise about 10^−3^ to 10^−4^ of the population according to NMR relaxation dispersion experiments, and this frequency falls within the range of translational errors ^53^. Based on this, we hypothesized that G:U mispairings should be distributed among the 1^st^ and 2^nd^ positions, but not the 3^rd^, and, since they are not amino acid but rather nucleotide-dependent, they should have no amino acid bias. Indeed, G:U mispairings partitioned exclusively, and at similar levels, between the first two codon-anticodon positions for both organisms (Fig. 1G, S2D). Furthermore, G:U events are present across a diverse range of codons and amino acids (Fig. 1F, S2C), consistent with the idea that tautomeric events depend only on the nucleobase chemistry and not the codon family.

Since G:U with one nucleobase in the *enol* tautomer result in stable G:C-like interactions, we then evaluated whether GC codon:anticodon content, a proxy for stability, correlated with the observed range of errors. In yeast, codons with higher than median error frequencies have increased GC content, whereas those with lower error frequencies are AU rich (Fig. S2E). GC and AU rich codons have a measured difference of 2-3kcal/mol ^54^, based on standard base-pair thermodynamics, which Grosjean and Westhof used as basis for an energy-based organization of the genetic code ^55^ (Fig. 1H). We grouped codon translation error frequencies according to their corresponding energy-based category. In yeast, we found a stark separation, where codons with strong codon-anticodon interactions have significantly higher error frequencies than those with weaker interactions, while intermediate interactions clearly bridge both extremes (Fig. 1I). There is a finer granularity to this general partitioning. For example, there are 4 strong Arginine codons and 2 intermediate ones: the strong GGC(Arg) triplet has an error frequency an order of magnitude higher than the intermediate AGA(Arg) codon. Similarly, Leucine codons are either intermediate (4 codons) or weak (2 codons): error frequency for the intermediate CUA(Leu) codon is about 5 times higher than frequency of the weak UUG(Leu) triplet, highlighting how both GC content and GC position are important. These results suggest that as codon-anticodon interactions get more stable based on thermodynamics, then their subsequent mispaired base-pairing configurations in near-cognate selection would also be more stable, leading to higher error frequencies in any organism. However, a similar analysis of the mouse estimates displayed an inverse trend, except for those codons with 100% GC content (Fig. 1J, S2F), suggesting that factors beyond pairing thermodynamics/kinetics play an essential role in determining fidelity.

### tRNA levels per codon determine translation fidelity

To explore the opposing relationships between codon-anticodon stability and error frequencies observed for yeast and mouse, we analyzed the levels of the codon (codon usage) and anticodon (tRNA isoacceptor) components in both organisms (Fig. 2A). We binned codon usage according to the described energy-based classification. In yeast, we observed a marked negative correlation between codon usage and codon stability (Fig. 2B left panel), whereas the mouse translatome exhibited a uniform codon usage across all stability classes (Fig. 2D left panel). Then, we measured the aminoacylated tRNA pool composition in both organisms using modification-induced misincorporation tRNA sequencing ^56^. While no major differences were observed for cognate tRNA levels between both organisms (Fig. S3A,B), near-cognate tRNA levels displayed different trends, with yeast levels being uniform across energy-based codon categories (Fig. 2B center) and mouse near-cognate tRNA levels decreasing as stability increases (Fig. 2D center). We integrated both observations by computing normalized per codon near-cognate tRNA levels, as the quotient of near-cognate tRNA fraction over codon frequency in the translatome, and binned them according to the energy-based classification (Fig. 2B right, 2D right). The resulting trends for both organisms recapitulate the opposing directionalities as the error frequencies binned by codon-anticodon stability (Fig. 1I-J). We directly compared the normalized per codon near-cognate tRNA levels against the estimated translation error frequencies, resulting in a significant positive correlation for both organisms (R=0.72 yeast, R=0.69 mouse; Fig. 2C,E). These analyses suggest that per codon near-cognate tRNA levels are a major driver for misincorporation, able to dominate the contribution that codon:anticodon thermodynamics alone has on fidelity. Of note, in yeast, five codons have unusually high error rates but, as a group, they also exhibit a positive correlation with their per codon near-cognate tRNA levels (R=0.94, Fig. 2C, S3C). The mouse data set has three characteristic outliers, two of which are phenylalanine codons with exceedingly high error rates and the lowest per codon near-cognate tRNA levels. We cannot rule out that these outliers present high error rates due to additional factors beyond tRNA concentrations, such as heavy misacylation. Thus, if codon usage remains constant, perturbations in near-cognate tRNA levels should also perturb error frequencies at the corresponding codons.

**Figure 2.**
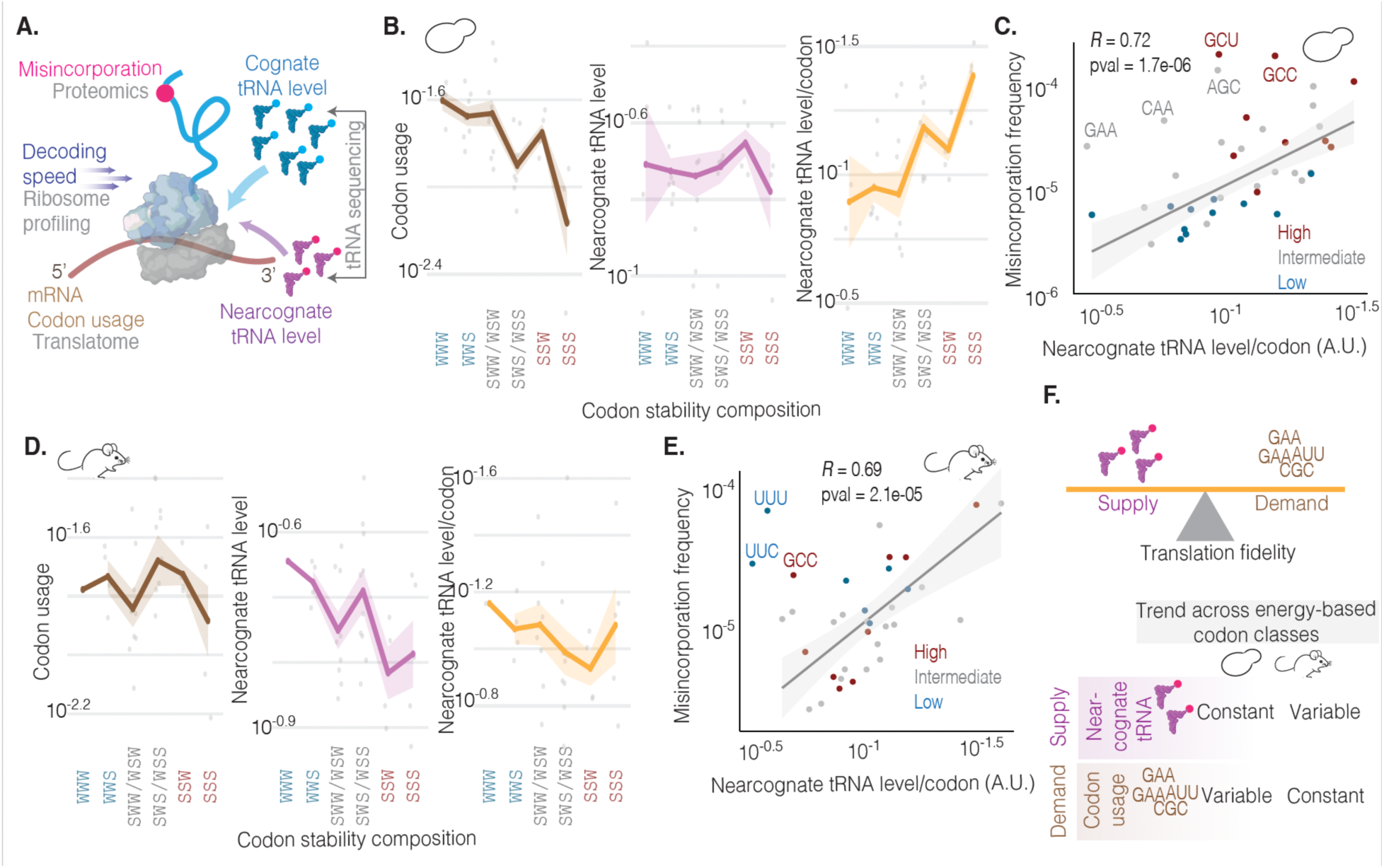
Mechanistic bases of amino acid misincorporation. A) Molecular factors known to play a role in protein translation fidelity. B) Panels depicting, for the yeast codons with measured misincorporation frequencies, codon usage (left), near-cognate tRNA level (middle), and the resulting near-cognate level over codon-usage values (right), binned using the energy-based codon classification. Solid line corresponds to the mean and shaded area to the standard error of the mean (SEM). C) Correlation plot between yeast codon misincorporation frequencies and their corresponding near-cognate tRNA levels over codon usage values; correlation by Pearson’s R. D) Panels depicting, for the mouse codons with measured misincorporation frequencies, codon usage (left), near-cognate tRNA level (middle), and the resulting near-cognate level over codon-usage values (right), binned using the energy-based codon classification. Solid line corresponds to the mean and shaded area to the SEM. E) Correlation plot between mouse codon misincorporation frequencies and their corresponding near-cognate tRNA levels over codon usage values; correlation by Pearson’s R. F) Schematic depicting the observed relationship between near-cognate tRNA levels, codon usage, and translation fidelity (top). Summary of the observed general trends for near-cognate tRNA levels and codon usage based on the energy-based codon binning analysis.

To test this hypothesis, we overexpressed tRNA^Arg^_CCU_ in *S. cerevisiae* (whose genome harbors a single copy of this gene) by ectopic expression via a plasmid. This resulted in a significant increase in near-cognate tRNA levels respect to cognate levels for the codons AGA(Arg) and UGG(Trp), as measured by tRNA sequencing (Fig. S4A), making UGG a candidate to map W>R mutations caused by increased near-cognate tRNA levels. W>R substitutions introduce new trypsin sites; therefore, we searched for those potential new tryptic peptides. We observed a drastic increase in W>R substitution frequency, 91%, upon tRNA^Arg^_CCU_ overexpression (Fig. S4B), supporting this hypothesis. Together, our results demonstrate that a balance between codon usage and near-cognate tRNA abundance governs translation fidelity, despite the intrinsic codon biases of an organism (Fig. 2F).

The error frequencies discussed so far (total error frequencies) encompass both near- and non-cognate misincorporations, with the former being dominant in both studied organisms (72% of total errors in yeast, 59% in mouse), consistent with previous reports ^2,3^. Near-cognate error frequencies correlate with near-cognate tRNA levels in a yeast and mouse (R=0.69 yeast, R=0.78 mouse; Fig. S5A,B), which is expected since they are the direct consequence of near-cognate tRNA misincorporation. Unexpectedly, non-cognate error frequencies also displayed a positive, albeit weaker, correlation with near-cognate tRNA levels (R=0.53 yeast, R=0.56 mouse; Fig. S5C,D). Non-cognate errors can originate through the reading of a codon by either a misacylated cognate or near-cognate tRNA, or by a properly aminoacylated non-cognate tRNA (two or more mispaired nucleotides). The ribosome efficiently rejects the latter during the initial tRNA sampling and kinetic proofreading ^31,38^, leaving misacylation as a more likely scenario. The observed correlation between near-cognate tRNA levels and non-cognate error frequencies suggests that misacylation of the near-cognate tRNA pool, due to misrecognition of tRNAs by aaRS, might generate non-cognate error. To explore this, we rationalized misacylation of cognate tRNAs which, for most codons, are less abundant than near-cognates, should also contribute to non-cognate errors. Thus, we compared per codon cognate-plus-near-cognate tRNAs against non-cognate error frequencies, resulting in an improved correlation respect to near-cognate tRNAs alone (Fig. S5E,F). This effect is more pronounced in yeast than mouse, because in yeast the covariation between cognate and near-cognate levels is lower (R=0.33, Fig. S5G) than in mouse (R=0.59, Fig. S5H), allowing both variables to exert stronger independent effects.

### Structural and functional implications of protein translation fidelity

Having mapped naturally occurring mutations in yeast and mouse, we next leveraged our dataset to understand the structural and functional implications of translational errors on the eukaryotic proteome, and how they contribute to phenotypic variation. Translation is a vectorial process during which the chemistry of short peptide stretches dominate the early stages of protein folding ^57–59^. Therefore, we started by analyzing the effect the identified mutations would exert on their local peptide context. Metagene analyses of solubility values for yeast misincorporation variants reveal near-cognate variants affect soluble regions, whereas non-cognate ones localize to insoluble peptides. Despite this difference, both types of mutations caused negligible change on local solubility compared to the wild-type (WT, Fig. 3B). A similar analysis on mouse misincorporation variants shows that near and non-cognate mutations map to local dips in solubility, more pronounced for non-cognate ones; while near-cognate variants have almost no effect in solubility, non-cognate mutations slightly increased it (Fig. S6A). Hence, naturally occurring translation errors, especially near-cognate ones (which constitute the majority), do not greatly impact solubility. This finding, on one hand speaks to the general organization of the genetic code (which has error suppressing qualities ^60^), and more specifically to the fact that near-cognate misincorporations more frequently involve mispairing at the first position of the codon whereas amino acid physiochemical properties are more heavily segregated by the second position of codons.

**Figure 3.**
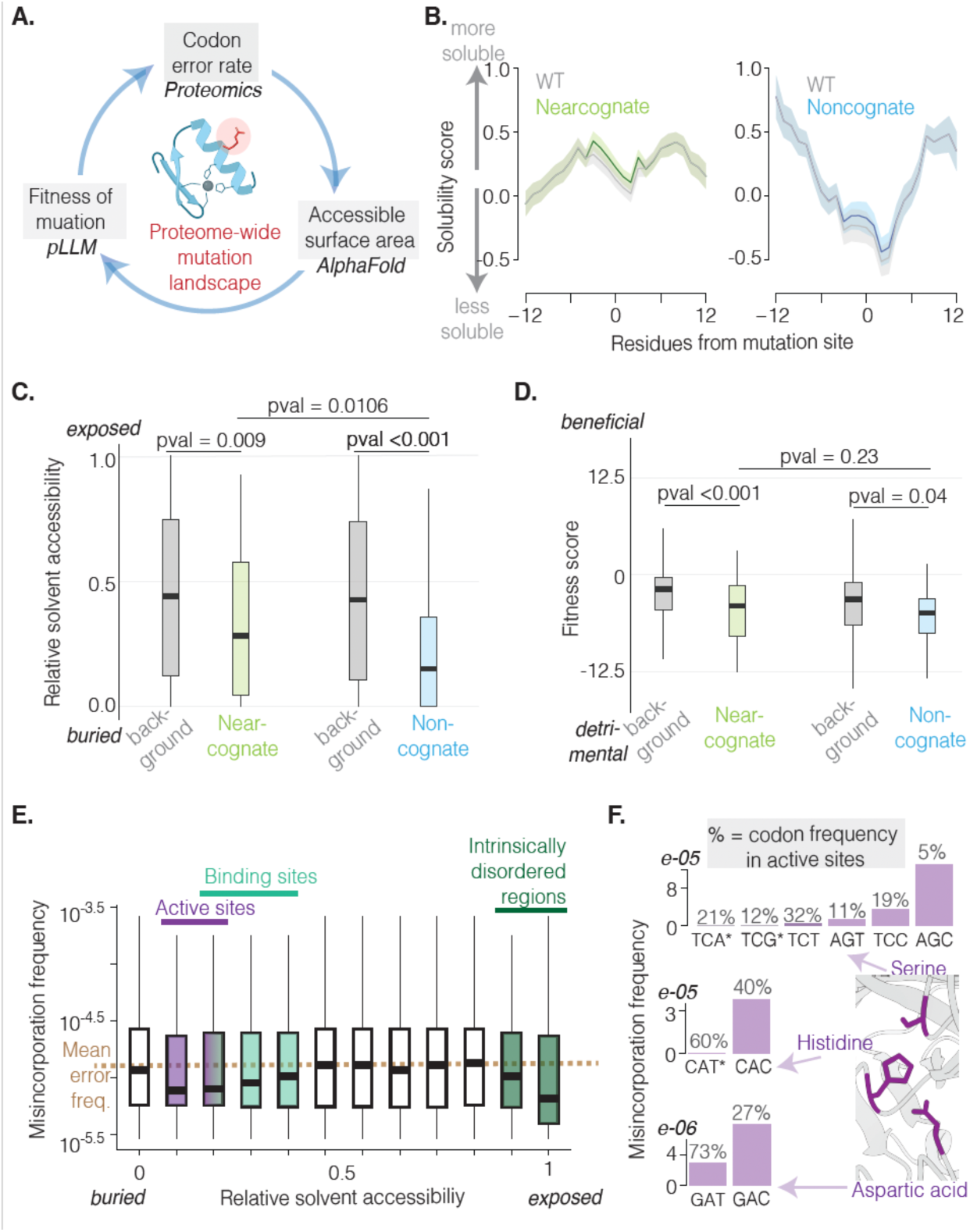
Structural and functional implications of protein translation errors in yeast. A) Components of the approach used to understand structural and functional implications of the mapped misincorporations. B) Sequence-based solubility values for all identified misincorporation variants (green – near-cognate, blue non-cognate), and their wild-type counterparts (grey), in their local peptidic context. Misincorporation position centered at position zero, x-axis, flanked by 12 residues at each side; mean (solid line) and SEM (shaded area). C-D) Box and whisker plots showing relative solvent accessibility values (RSAs, C) and fitness scores (D) for the identified misincorporation variants, compared to their respective background distributions (i.e. RSAs or fitness scores for all possible near- or non-cognate mutations in the proteome). P-value for near- and non-cognate comparisons according to Wilcoxon test; comparisons against background by bootstrapped Wasserstein distance. E) Misincorporation frequencies for each proteome RSA decile – extreme values (fully exposed, RSA = 0; fully buried, RSA = 1) are plotted separate from their corresponding deciles. Colored deciles correspond to the RSA ranges of functional elements. F) Codon misincorporation frequencies for synonymous codons encoding different catalytic residues. Percentages indicate prevalence at codons encoding catalytic sites. Asterisks indicate non-observed substitutions, thus frequencies are at most 1/coverage: * < 1e-05; ** < 2.3e-05; *** < 1.1e-05.

Next, we evaluated the spatial distribution of the identified mutations by mapping them onto their corresponding 3D AlphaFold structures ^61^ and computing their corresponding relative solvent accessibilities (RSA). In both organisms we found near-cognate misincorporation variants to be more solvent accessible than non-cognate ones (Fig. 3C, S6B). However, compared to all possible near- or non-cognate mutations in the yeast or mouse proteomes, substitutions occur, on average, at sites that are more buried than expected (Fig. 3C, S6B). To assess possible mutational effects, we estimated a mutational fitness score using a structure-aware protein large language model (pLLM), where a positive score indicates that a substitution is more favorable than the WT, from an evolution point of view, and negative scores signal otherwise ^62^. We computed fitness scores for all possible amino acid substitutions in both yeast and mouse proteomes, i.e. background distributions, and compared them against the scores of the identified substitutions. In both organisms, most identified misincorporation variants have negative scores, as do their respective background distributions, with non-cognate ones having scores more negative than expected from random sampling of the background distribution (Fig. 3D,S6C). Non-cognate substitutions are in general more negative than near-cognate ones (Fig. 3D,S6C). Importantly, our observations are not due to proteomics sampling bias, since the sequenced WT residues evenly sample the corresponding overall organismal amino acid substitution space (S6D,E).

We wondered if the observed non-random spatial distribution of errors could serve as a potential safeguarding mechanism for critical residues. Therefore, we explored whether there is an interplay between misincorporation, RSA, and protein functional sites. We annotated several functional classes (active site, binding site, IDR) according to how they distribute by RSA in the yeast proteome (Fig. S6F). Then, we calculated the error frequency distribution for various RSA bins and overlayed the annotated RSA ranges of the functional classes. This analysis reveals most annotated functional sites fall in RSA bins with a lower-than-average error rate (Fig. 3E). Zooming in in one of the functional classes, active sites, we asked if catalytic residues are preferentially encoded by low-error codons, as there should be selective pressure to guarantee their faithful decoding. In yeast, catalytic serine, histidine, and aspartic acid residues are preferentially encoded by the synonymous codon(s) with the lowest error rate(s) (Fig. 3F). The mouse data set exhibits the same trend for histidine and aspartic acid, but only one serine codon was quantified (Fig. S6G). Our analyses suggest that this non-random distribution of errors and use of low-error codons in crucial active sites are an important aspect of translation fidelity, in turn ensuring protein functionality.

### Protein error frequencies increase during yeast chronological aging as a result of tRNA pool remodeling

The protein error landscapes described so far result from balanced cellular and organismal proteostasis, a product of young cells growing in optimal conditions. However, as cells and organisms age, their proteostasis capacity declines ^63,64^. To test whether aging-related proteostasis disruption could produce aberrant translation error patterns and frequencies, we first studied chronological aging using *S. cerevisiae* as a model ^65^. Four time points were considered, one representing actively dividing cells, day 0 – corresponding to the yeast data discussed so far, and three (days 2, 4, 6) corresponding to a post-mitotic aging state, where starvation is pervasive (Fig. 4A). We measured protein error frequencies in the total proteome and compared them across timepoints. Total misincorporation frequencies drastically increased with age, with the error transition from mitotic to post-mitotic growth, day 0 to day 2, being the most pronounced (Fig. 4B). Separating those frequencies into their near- and non-cognate components reveals a similar upward trend for both classes, with near-cognate errors being dominant across all time points (Fig. S7A). Importantly, the increase in total misincorporation frequency stems from a common behavior by the individual codon error frequencies, as almost every codon measured across all four ages exhibits an increase in error frequency (Fig. S7B). Interestingly, SCR frequency did not exhibit a similar age-related increase (Fig. S7C).

**Figure 4.**
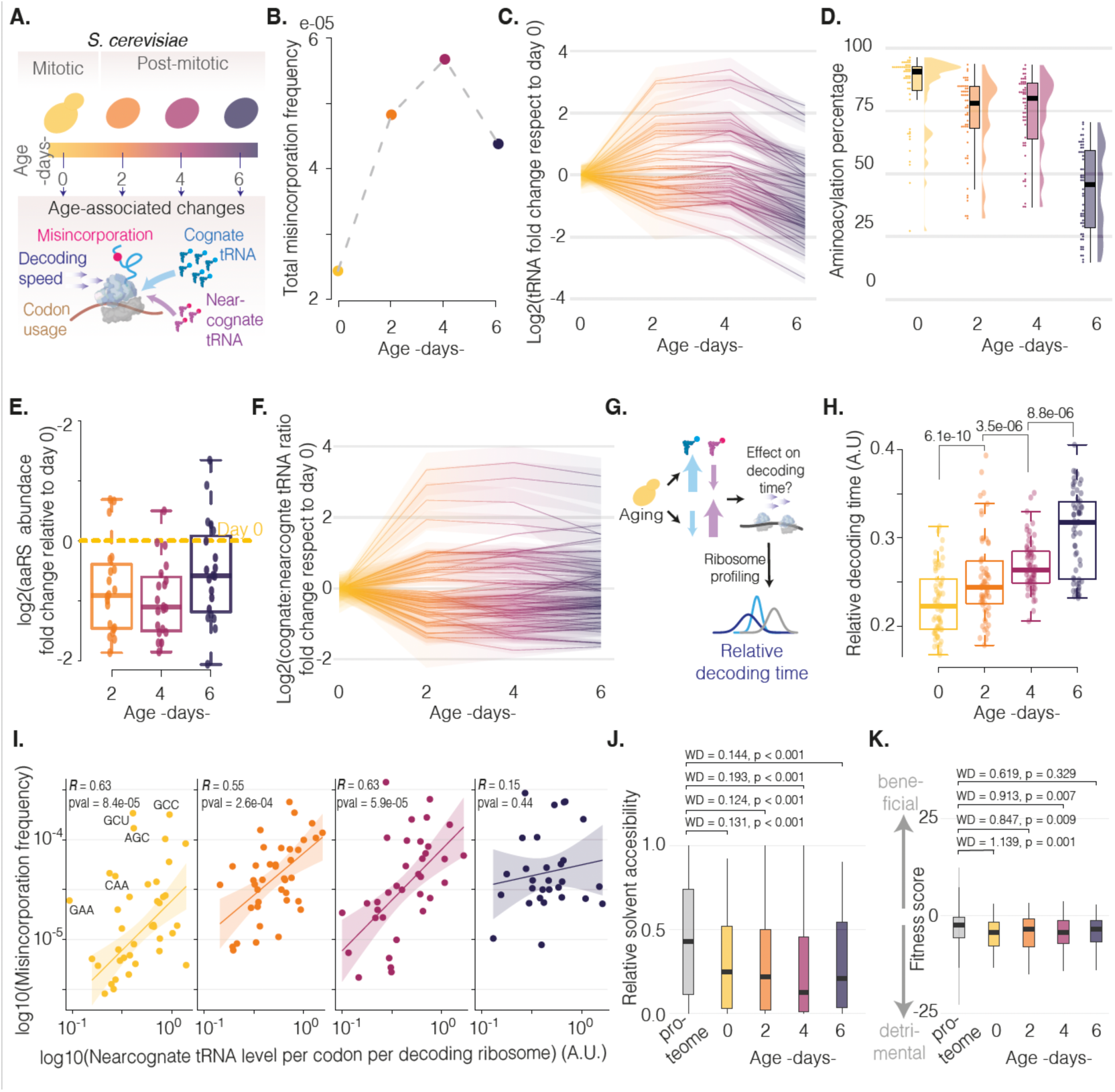
The age-dependent protein error landscape in *S. cerevisiae*. A) Time points from chronologically aged yeast cultures (n = 3) used to study translation fidelity during aging. Color scheme for each timepoint is consistently used in all other panels. B) Total amino acid misincorporation frequency as a function of yeast age. C) Fold change in aminoacylated tRNA isoacceptor level respect to day 0, each line represents a different isoacceptor. D) Distribution of tRNA isoacceptor aminoacylation percentages at each evaluated time point. E) Fold change in aminoacyl tRNA synthetase abundance respect to day 0, by label-free quantification of proteomics data. F) Fold change in cognate:near-cognate tRNA levels respect to day 0, each line corresponds to a different sense codon. G) Schematic representing the influence of tRNA pool remodeling on codon decoding time. H) Relative decoding times for each sense codon, estimated from ribosome profiling data, at each aging time point; p-values from Wilcoxon test. I) Codon misincorporation frequencies as a function of near-cognate tRNA level per codon per decoding ribosome, for every yeast age; correlation by Pearson’s R. J-K) Relative solvent accessibility values (J) and fitness scores (K) for the identified misincorporation variants at each time point, compared to the corresponding background distribution (RSA or fitness scores for all possible mutations); bootstrapped Wasserstein distance p values.

The position and type of codon:anticodon mispairings responsible for the age-related misincorporations greatly mimic the patterns observed at day 0, suggesting a shared mechanistic origin (Fig. S7D,E). At day 0 two important variables correlated with error frequency: codon-anticodon stability and near-cognate tRNA availability. However, the relationship between error frequency and stability no longer holds following day 2(Fig. S7F-H). If we assume that codon-anticodon stabilities in the mechanism of tRNA selection are constant with age, then near-cognate tRNA availability is a potential candidate to explain the observed age-associated increase in error frequency. To test this, we assessed the temporal dynamics of the tRNA pool by tRNA sequencing. We indeed observed a wide age-related variation in the aminoacylated tRNA pool (Fig. 4C), which stemmed from the combination of changes in total tRNA abundance (Fig. S7I) and an age-associated decrease in aminoacylation (Fig. 4D, S7J), linked to a downregulation of amino acyl tRNA synthetases (Fig. 4E). Next, we computed the resulting per codon cognate and near-cognate tRNA levels. With age, the normalized per codon ratio of cognate to near-cognate tRNAs decreased for two thirds of the codons (Fig. 4F).

Cognate tRNA levels are a major determinant of decoding speed ^31,38,44^. To observe whether age-associated changes in cognate tRNA levels could alter translation decoding times and misincorporation frequencies, we performed ribosome profiling (Ribo-seq) in yeast cells of all analyzed ages, in the presence of tigecycline (TIG) and cycloheximide (CHX) (Fig. 4G). The TIG/CHX cocktail captures ribosomes in the pre-accommodation state, yielding small ribosome footprints proportional to decoding time ^66^. We found an age-related, gradual increase in small ribosome footprints (Fig. S8A,B), indicating that codon decoding times slow down with age. To investigate if this effect is general or driven only by a few codons, we calculated relative individual codon decoding times using the ribosome profiling data. We found, at all older timepoints, the relative decoding times of most codons increase, with only a few decreasing (Fig. 4H, S8C-E). Accordingly, at all older timepoints, codons with decreased decoding time have higher cognate tRNA availability than codons with increased decoding times (Fig. S8F-H).

Our tRNA-seq and Ribo-seq analyses support the idea that age-related remodeling of the tRNA pool directly affects codon decoding speed, exposing codons to a higher time window of near-cognate tRNA sampling and therefore increasing the probability of misincorporation. To probe this, we investigated if the positive correlation between codon misincorporation frequencies and near-cognate tRNA levels per codon observed at day 0 (Fig. 2C) also holds in old cells. The resulting correlations were weak (Fig. S9A-C). We rationalized that this could be because the age-related tRNA pools change both near-cognate tRNA frequencies and decoding speed, i.e. the time of near-cognate tRNA sampling. If this was the case, including decoding speed in the calculations should restore the correlation. Indeed, by accounting for changes in decoding speed (i.e. tRNA levels per codon per decoding ribosome), the correlations with error frequencies were restored for days 2 and 4, but not day 6 (Fig. 4I), suggesting that by day 6 other factors, such as mutation bias stemming from error accumulation in the cell, or differential degradation patterns, influence the measured error frequencies. These analyses demonstrate that an age- (and starvation-) driven remodeling of the aminoacylated tRNA pool leads to increased error frequencies in the proteome of aging yeast cells.

We next sought to understand the possible functional and structural consequences of increased error frequencies in the aging proteome. We first mapped all observed misincorporation variants onto their corresponding 3D structures and analyzed their relative solvent accessibilities. We observed an age-associated migration of errors towards buried regions (Fig. 4J), mainly driven by near-cognate variants, as non-cognate substitutions were buried deeper in the non-aged experiments (Fig. S9D,E). Next, to assess the possible functional impact of the observed misincorporation variants, we compared their fitness score distributions across all ages. Almost all observed variants carry a negative fitness score (Fig. 4K), but their distributions did not follow the same age-related decline as the RSA distributions. However, by partitioning mutation variants into cognate and near-cognate, we found that non-cognate ones score more negatively with age (Fig. S9F,G). This reflects the fact that, at day 0, near- and non-cognate fitness scores distributions are not statistically different, whereas from day 2 non-cognate errors are more negative than near-cognates (Fig. S9H). Importantly, these patterns are not a sampling artifact given how the fitness score search space available through our proteomics data evenly samples the entire yeast mutational space (Fig. S9I), underscoring the depth of our data.

### Protein error frequency is organ- and age-dependent in mouse

To investigate whether age-related change in protein error frequency could be a more general phenomenon across eukaryotes, we analyzed samples from aging mice. We compared liver, which has a high protein synthesis rate and mitotic capacity ^67^ despite its quiescent state, against heart, predominantly post-mitotic and with lower synthesis rates ^67^, coming from mice 3-, 18-, and 30-month-old (Fig. 5A). First, we estimated their protein error frequencies using our mass spectrometry approach. We found both the misincorporation and the SCR frequencies in the heart to be higher than in liver, even at 3 months of age (Fig. 5B, S10A). Furthermore, misincorporation frequencies increase with age in the heart, similar to aging yeast, but remain steadier in liver samples (Fig. 5B). These trends are similar when analyzing near- and non-cognate error frequencies separately (Fig. S10B,C). Similar to misincorporation with age SCR levels increased in the heart and remained lower in the liver (Fig. S10A). Our measurements reveal that translation fidelity is not only age-, but also organ-dependent.

**Figure 5.**
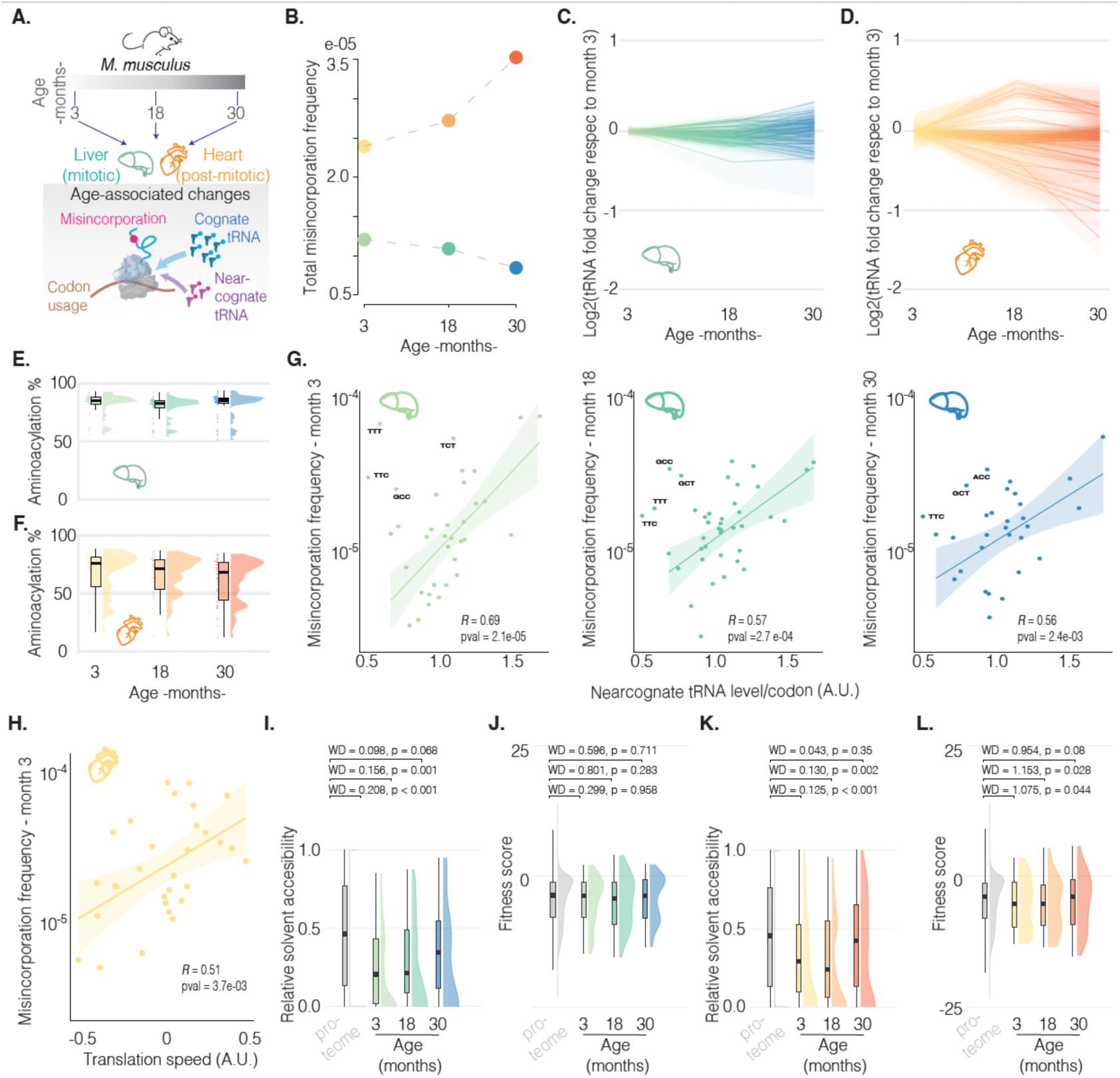
Organ- and age-dependent dynamics of protein errors in aging mice. A) Organs from mice (n = 3) of different ages used to study translation fidelity during mouse aging. B) Total amino acid misincorporation frequency, as a function of age, for mouse liver (green-blue) and heart (yellow-red). C-D) Fold change in aminoacylated tRNA isoacceptor level compared to month 3 for liver (C) and heart (D); each line represents a different isoacceptor. E-F) Distribution of tRNA isoacceptor aminoacylation percentages at each evaluated age; (E) lliver, (F) heart. G) Liver codon misincorporation frequencies as a function of near-cognate tRNA level per codon, for every evaluated age; correlation by Pearson’s R. H) Heart (month 3) codon misincorporation frequencies as a function of codon translation speed; correlation by Pearson’s R. I,J) Relative solvent accessibility values (I) and fitness scores (J) for the identified liver misincorporation variants at each age, compared to the corresponding background distribution (RSA or fitness scores for all possible mutations); bootstrapped Wasserstein distance p values.. K,L) Relative solvent accessibility values (K) and fitness scores (L) for the identified heart misincorporation variants at each age, compared to the corresponding background distribution; bootstrapped Wasserstein distance p values.

Considering the measured liver and heart error frequencies together with our yeast findings, we hypothesized that aging liver samples should exhibit minor perturbations in tRNA pool composition, whereas heart samples should display a more prominent age-related variation. To test this, we carried out tRNA sequencing across all time points, for both organs. In the liver, consistent with our hypothesis, we observed little age-related variation in the aminoacylated isoacceptor tRNA pool (Fig. 5C). However, the aminoacylated isodecoder tRNA pool had greater age-related variation (Fig. S10D), indicating that, during aging, opposing changes in isodecoders balance the final isoacceptor tRNA levels, similar to findings during cell differentiation and within-tissue comparisons ^68,69^. We found these trends to be driven by changes in tRNA abundance rather than aminoacylation, as the latter had virtually no change with age (Fig. 5E, S10E). On the other hand, heart-derived samples exhibited detectable age-related changes in the aminoacylated isoacceptor tRNA pool (Fig. 5D), though these are smaller changes than we observed in yeast, likely because in this case aging is not coupled to starvation like it is in the yeast chronological aging model. Unlike the liver, heart isodecoders vary in a similar manner to isoacceptors (Fig. S10F) and thus, in this case, isoacceptor changes are synergistic rather than opposing. Interestingly, unlike the liver, aminoacylation levels vary with age in the heart (Fig. 5F, S10G), and this plays a role in accentuating changes in total aminoacylated tRNA abundance.

Based on the steady misincorportion frequencies and aminoacylated tRNA levels in the aging liver, we hypothesized that the positive correlation between near-cognate tRNA levels per codon and error frequencies we observed earlier (Fig. 2E) should hold with age. We computed near-cognate tRNA levels for all ages and compared them against the corresponding error frequencies, resulting in similar positive correlations across all ages (Fig. 5G), further validating the fundamental role of the tRNA pool in translation fidelity. However, when performing a similar analysis for heart-derived samples, we saw no correlation between near-cognate tRNA levels and error frequencies (Fig. S11A-C). This makes sense based on our yeast results, where post-mitotic timepoints also had poor correlations unless ribosome decoding speed was accounted for. We tested this by estimating codon translation speeds using public ribosome profiling data from 10-week-old hearts of the same mouse strain ^70^; they do not reflect actual decoding times as our TGI/CHX yeast data because the footprints correspond to ribosomes across the entire elongation cycle. Nevertheless, we found a positive correlation between the estimated translation speeds and error frequencies (Fig. 5H), on par with liver correlations, suggesting that cardiomyocytes in post-mitotic state do follow a similar behavior as chronologically aged post-mitotic yeast.

Finally, we evaluated the spatial distribution and possible functional impact of the observed mouse mutations. When mapped onto their 3D structures, RSA distributions for mutations from the first two time points of both heart and liver show a statistically significant bias towards buried regions (Fig. 5I, K), resembling the observations from aging yeast. At month 30, in both organs, mutations follow a more uniform distribution, more significantly so for heart (Fig. 5I, K). When comparing their fitness scores, we found that all liver mutations, regardless of age, are distributed following the background distribution, i.e. score distribution of all possible mutations (Fig. 5J). In short, the mutational fitness burden in the liver does not change with age. On the other hand, fitness scores for heart mutations resemble the trend observed in aging yeast, where mutations tend to be more detrimental than expected from the background distribution (Fig. 5L). Separating heart mutation fitness scores into near- and non-cognates reveals a divergent behavior: with age, near-cognate distributions become more on par with the background, whereas noncognate ones become more negative (Fig. S11D,E), also resembling our yeast results (Fig. S9G,H). As a consequence, with age, non-cognate mutations impose a significantly higher fitness burden respect to near-cognate ones (Fig. S11F). These trends were not observed in a similar analysis of the liver mutations (Fig. S11G-I) and, importantly, they are not the product of sequencing bias since the distributions of unmodified residues properly sample the proteome (Fig. S11J,K).

## Discussion

Although advances in next generation sequencing (NGS) have driven genome-wide interrogation of nucleic acids, proteomics has lagged behind NGS in terms of sequencing depth, limiting the ability to track global translational errors. Previously, proteome-wide misincorporation measurements in eukaryotes have only been achieved by collating most of the available public proteomics data sets ^3^. Here we demonstrate that shotgun proteomics can routinely interrogate single amino acid mutations in a proteome-wide fashion, even for mammalian proteomes. We achieve unbiased estimates for about two thirds of the genetic code thanks to even sampling of the proteomic space. We envision that missing codons can be quantified by heavy off-line sample fractionation, and/or use of equipment specialized in capturing rare ions. Since our error frequency estimates are based on spectral counting and, despite de achieved sequencing depth, we acknowledge that the reported frequencies are not absolute error frequencies and might be limited by rare ion under sampling. Nevertheless, our current measurements allow us to probe two fundamental questions: what are the causes and consequences of protein synthesis errors? Do such errors increase with age?

Across the tree of life, the translation machinery is among the most conserved biological systems ^71^. Therefore, we first hypothesized fundamental conserved mechanisms for translational errors. Tautomerization of uracil nucleobases during codon-anticodon pairings has been proposed as a major source of misincorporation, especially formation of tautomeric G-U pairs that mimic Watson-Crick geometry in the conserved decoding center of the ribosome ^50,72^. This keto-enol equilibrium is, fundamentally, taxa agnostic. Indeed, our data suggests G:U*/G*:U tautomeric pairs (particularly at position 1 of the codon) are the most likely source of misincorporations in both yeast and mouse, correlating with known thermodynamic and kinetic aspects of codon-anticodon pairing ^55^. In yeast, codons prone to forming stable codon-anticodon interactions (G-C rich) have higher error rates than less stable pairings. Upon misreading, preserving stable interactions is more likely in G-C rich pairings than in A-U counterparts, e.g. GGG misreading would preserve 2 G:C interactions and could potentially have a G:U* tautomer mispairing. Thus, there is an intrinsic error frequency trend based on simple codon-anticodon thermodynamics, with higher stability interactions having increased stability of near-cognate interaction leading to misincorporations. To further understand this interplay, a more detailed understanding of how tRNA modifications affect codon-anticodon stability and tRNA selection is needed, along with a careful consideration of variables such as organismal temperature.

Alongside codon-anticodon stability, tRNA substrate concentrations contribute to translational fidelity. During each round of translation elongation, all aminoacyl-tRNAs in ternary complex with eEF-1A (EF-Tu in bacteria) and GTP, sample the codon in the ribosomal decoding center in a concentration-dependent manner, with kinetic steps and coupled GTP hydrolysis leading to greatly preferred selection for cognate over near-cognate tRNAs ^31^. Our yeast analyses show that high per-codon near-cognate tRNA levels result in high error rates. In yeast, these two factors, codon-anticodon stability and tRNA concentration, have a synergistic effect: codons with higher per codon near-cognate tRNA availability are also codons with higher codon-anticodon stability, making their error frequencies the highest. However, that synergy is reversed in mouse, where the codons with higher near-cognate tRNA availability are those with lower codon-anticodon stabilities. Here the tRNA pool component of error frequency competes with the contribution of codon-anticodon stability and, in fact, dominates the general trend. The competing contributions of codon-anticodon stability and near-cognate tRNA concentration might be a reason for the lower total error frequency observed in mouse with respect to yeast.

Although near-cognate tRNAs have rapid initial and subsequent dissociation rates from the ribosome, non-cognate tRNAs dissociate almost immediately ^38^. Thus, it is not surprising that near-cognate substitutions are more frequent in both yeast and mouse, consistent with previous data from bacteria and yeast ^2,3^. What is surprising, instead, is that, despite their efficient initial rejection, non-cognate errors are 30-40% prevalent in our data. This suggests these residues were not delivered by non-cognate tRNAs, but by misacylated cognate or near-cognate tRNAs. Our data supports this mis-acylation model, given how well non-cognate error frequencies correlate with per codon cognate-plus-near-cognate tRNA levels. This highlights the question of how aminoacyl tRNA synthetases (aaRS) have co-evolved with translation speed and the tRNA pool to mitigate non-cognate errors.

In yeast, amino acid residues on the surface (high RSA) typically evolve at higher rates than core residues ^73,74^. Recent deep mutational scanning data of human domains also found protein core regions tend to be less tolerant to missense mutations ^75^. In other words, genetic mutations causing missense substitutions are more detrimental at buried sites. However, most of our mapped misincorporation variants do not occur at exposed regions, and, in fact, they are more buried than one would expect by random sampling of all possible misincorporations. Considering this unexpected RSA behavior, we expected our variants to have negative fitness effects, especially non-cognate substitutions since they tend to occur at even more buried sites than near-cognate substituions. On the other hand, residues in the protein core tend to be more involved in binding and catalysis, which means that misincorporation may be utilized to access phenotypic variation not as easily accessed by standard genetic mutation. Our results show that protein translation imposes a fitness burden that is not the smallest it could be, further supporting the idea of a trade-off between translation speed and fidelity.

In various model organisms, a drug- or genotype-induced decrease in translation fidelity shortens lifespan, whereas an increase extends it ^25,76,77^. At the core of these observations lingers a fundamental question, largely unanswered: do translation errors increase with age? Our data provide not only an answer, but also an important mechanistic lesson: elements regulating error frequencies, e.g. tRNA pool and ribosome decoding speed, change with age in an organ-specific manner and, thus, the answer can be yes, like in the heart, or no, as in the liver. Our results agree with recently documented age- and organ-dependent changes in stop codon readthrough frequencies using reporters ^78^ and provide insights to recently observed tRNA changes in cerebral cortex of aging killifish ^79^. The functional bases of the observed organ-specific behavior are unknown: are they related to protein synthesis capacity? The liver has a greater protein synthesis output than the heart, therefore lower fidelity would be more detrimental; or is it because cell cycle state compromises the ability to reset the tRNA pool? Hepatocytes, although quiescent, have mitotic capacity, and share traits with dividing yeast, whereas cardiomyocytes are post-mitotic cells and resemble patterns seen in chronologically aged yeast.

In liver, where we observe a constant tRNA pool composition and low error frequencies during aging, misincorporation events evenly sample the mutational space across all ages. Accordingly, a tight regulation of the tRNA pool seems to guarantee a manageable mutational burden by means of constant, if not low, error frequencies. Contrary to liver, in heart and yeast, even at the early time points, mutations are more detrimental than random and, in fact, non-cognate mutations worsen with age, at the same time that error frequencies increase and tRNA levels change. The initial, young configuration of the tRNA pool might define an accessible mutational space with an associated mutational fitness burden the cell can afford to meet its protein synthesis needs. The caveat of this initial tRNA pool configuration is that its dysregulated state is more likely to access detrimental mutations. Mitotic cells, such as our day 0 yeast, could have a frequent opportunity to reset the tRNA pool, avoiding risk of dysregulation. However, post-mitotic ones, like in our heart samples, might not be afforded that opportunity, making long-term tRNA pool regulation a challenging task. In rats, aging results in diminished protein synthesis rate in the heart, while rates in the liver barely change ^80^. Similarly, protein synthesis goes down in aging yeast, as seen by polysome profiles ^81^. Lowering the protein demand with age might serve as a mechanism to counterbalance high error frequencies.

The “life on the edge of solubility” hypothesis, shown to be a proteome-wide phenomenon, states that proteins are expressed close to their solubility limits ^82,83^. Aging tilts this balance towards aggregation ^83^, which has clear implications for age-associated proteinopathies. Our work opens a new avenue to understand the fundamental forces affecting the aging proteome and, hopefully, ways to restore its lost balance.

## Acknowledgments

We are grateful to members of Puglisi labs for helpful discussions and feedback. We thank Dr. Madhav Mantri, Douglas Henze, Dr. Andy P. Tsai, Dr. Ian H. Guldner for their assistance with mouse sample collection. We thank the Frydman lab for sharing the *S. cerevisiae tRNA^Arg^_CCU_* overexpression strain, originally from the Pilpel lab.

## Funding

National Institutes of Health grant GM145306 (JDP)

National Institutes of Health grant AG064690 (JDP)

Chan Zuckerberg Biohub Investigator Award (JDP)

National Institute of General Medical Sciences DP2-GM140926 (SDF)

Sloan Fellowship (SDF)

Norn Group, Hevolution Foundation and Rosenkranz Foundation - Longevity Impetus Grant (SDF)

National Institutes of Health grant T32-GM135131 (NBW)

National Institutes of Health grant T32GM080189 (HET)

National Institutes of Health grant T32AG027668 (HET)

## Author contributions

MAR and JDP. conceived and designed the study; MAR performed all experiments; MAR analyzed the data and wrote the software; CP, JL, HET, NBW sample preparation and processing with guidance from SF and SQ; MAR writing original draft; JDP writing, reviewing & editing; CP, JL, HET, NBW SF, SQ manuscript revisions; JDP supervision and funding acquisition.

## Competing interests

Authors declare that they have no competing interests.

## Notes

### Competing Interest Statement

The authors have declared no competing interest.

## References

1. Ke, Z. et al. Translation fidelity coevolves with longevity. Aging Cell 16, 988–993 (2017).

2. Mordret, E. et al. Systematic Detection of Amino Acid Substitutions in Proteomes Reveals Mechanistic Basis of Ribosome Errors and Selection for Translation Fidelity. Molecular Cell 75, 427–441.e5 (2019).

3. Landerer, C., Poehls, J. & Toth-Petroczy, A. Fitness Effects of Phenotypic Mutations at Proteome-Scale Reveal Optimality of Translation Machinery. Mol Biol Evol 41, msae048 (2024).

4. Tsour, S. et al. Alternate RNA decoding results in stable and abundant proteins in mammals. Nature 656, 506–515 (2026).

5. Tretyachenko, V. et al. Encoded and non-genetic protein variants expand human functional proteome. Nature 1–3 (2026) doi:10.1038/s41586-026-11124-z.

6. Loftfield, R. B. & Vanderjagt, D. The frequency of errors in protein biosynthesis. Biochem J 128, 1353–1356 (1972).

7. Edelmann, P. & Gallant, J. Mistranslation in E. coli. Cell 10, 131–137 (1977).

8. Parker, J. Errors and alternatives in reading the universal genetic code. Microbiol Rev 53, 273–298 (1989).

9. Kramer, E. B. & Farabaugh, P. J. The frequency of translational misreading errors in E. coli is largely determined by tRNA competition. RNA 13, 87–96 (2007).

10. Kramer, E. B., Vallabhaneni, H., Mayer, L. M. & Farabaugh, P. J. A comprehensive analysis of translational missense errors in the yeast Saccharomyces cerevisiae. RNA 16, 1797–1808 (2010).

11. Thompson, R. C. & Karim, A. M. The accuracy of protein biosynthesis is limited by its speed: high fidelity selection by ribosomes of aminoacyl-tRNA ternary complexes containing GTP[gamma S]. Proc Natl Acad Sci U S A 79, 4922–4926 (1982).

12. Ruusala, T., Andersson, D., Ehrenberg, M. & Kurland, C. G. Hyper-accurate ribosomes inhibit growth. EMBO J 3, 2575–2580 (1984).

13. Wohlgemuth, I., Pohl, C. & Rodnina, M. V. Optimization of speed and accuracy of decoding in translation. The EMBO Journal https://doi.org/10.1038/emboj.2010.229 (2010) doi:10.1038/emboj.2010.229.

14. Johansson, M., Zhang, J. & Ehrenberg, M. Genetic code translation displays a linear trade-off between efficiency and accuracy of tRNA selection. Proc Natl Acad Sci U S A 109, 131– 136 (2012).

15. Xie, J. et al. Regulation of the Elongation Phase of Protein Synthesis Enhances Translation Accuracy and Modulates Lifespan. Current Biology 29, 737–749.e5 (2019).

16. Johansson, M., Lovmar, M. & Ehrenberg, M. Rate and accuracy of bacterial protein synthesis revisited. Curr Opin Microbiol 11, 141–147 (2008).

17. Jürgenstein, K. et al. Variance in translational fidelity of different bacterial species is affected by pseudouridines in the tRNA anticodon stem-loop. RNA Biol 19, 1050–1058.

18. Davies, J., Gorini, L. & Davis, B. D. Misreading of RNA codewords induced by aminoglycoside antibiotics. Mol Pharmacol 1, 93–106 (1965).

19. Davies, J. & Davis, B. D. Misreading of ribonucleic acid code words induced by aminoglycoside antibiotics. The effect of drug concentration. J Biol Chem 243, 3312–3316 (1968).

20. Pape, T., Wintermeyer, W. & Rodnina, M. V. Conformational switch in the decoding region of 16S rRNA during aminoacyl-tRNA selection on the ribosome. Nat Struct Biol 7, 104– 107 (2000).

21. Rosset, R. & Gorini, L. A ribosomal ambiguity mutation. J Mol Biol 39, 95–112 (1969).

22. Gorini, L. & Kataja, E. PHENOTYPIC REPAIR BY STREPTOMYCIN OF DEFECTIVE GENOTYPES IN E. COLI. Proc Natl Acad Sci U S A 51, 487–493 (1964).

23. Ozaki, M., Mizushima, S. & Nomura, M. Identification and functional characterization of the protein controlled by the streptomycin-resistant locus in E. coli. Nature 222, 333–339 (1969).

24. Martinez-Miguel, V. E. et al. Increased fidelity of protein synthesis extends lifespan. Cell Metab 33, 2288–2300.e12 (2021).

25. von der Haar, T., et al. The control of translational accuracy is a determinant of healthy ageing in yeast. Open Biol 7, 160291 (2017).

26. Crick, F. H. C., Barnett, L., Brenner, S. & Watts-Tobin, R. J. General Nature of the Genetic Code for Proteins. Nature 192, 1227–1232 (1961).

27. Nirenberg, M. W. & Matthaei, J. H. The dependence of cell-free protein synthesis in E. coli upon naturally occurring or synthetic polyribonucleotides. Proc Natl Acad Sci U S A 47, 1588–1602 (1961).

28. Martin, R. G., Matthaei, J. H., Jones, O. W. & Nirenberg, M. W. Ribonucleotide composition of the genetic code. Biochem Biophys Res Commun 6, 410–414 (1962).

29. Fujishima, K. & Kanai, A. tRNA gene diversity in the three domains of life. Front Genet 5, 142 (2014).

30. Varenne, S., Buc, J., Lloubes, R. & Lazdunski, C. Translation is a non-uniform process. Effect of tRNA availability on the rate of elongation of nascent polypeptide chains. J Mol Biol 180, 549–576 (1984).

31. Rodnina, M. V. & Wintermeyer, W. Fidelity of aminoacyl-tRNA selection on the ribosome: kinetic and structural mechanisms. Annu Rev Biochem 70, 415–435 (2001).

32. Rodnina, M. V. & Wintermeyer, W. Ribosome fidelity: tRNA discrimination, proofreading and induced fit. Trends Biochem Sci 26, 124–130 (2001).

33. Blanchard, S. C., Gonzalez, R. L., Kim, H. D., Chu, S. & Puglisi, J. D. tRNA selection and kinetic proofreading in translation. Nat Struct Mol Biol 11, 1008–1014 (2004).

34. Ogle, J. M., Murphy, F. V., Tarry, M. J. & Ramakrishnan, V. Selection of tRNA by the ribosome requires a transition from an open to a closed form. Cell 111, 721–732 (2002).

35. Ogle, J. M. & Ramakrishnan, V. Structural insights into translational fidelity. Annu Rev Biochem 74, 129–177 (2005).

36. Hopfield, J. J. Kinetic proofreading: a new mechanism for reducing errors in biosynthetic processes requiring high specificity. Proc Natl Acad Sci U S A 71, 4135–4139 (1974).

37. Ninio, J. Kinetic amplification of enzyme discrimination. Biochimie 57, 587–595 (1975).

38. Gromadski, K. B. & Rodnina, M. V. Kinetic Determinants of High-Fidelity tRNA Discrimination on the Ribosome. Molecular Cell 13, 191–200 (2004).

39. Lee, T.-H., Blanchard, S. C., Kim, H. D., Puglisi, J. D. & Chu, S. The role of fluctuations in tRNA selection by the ribosome. Proceedings of the National Academy of Sciences 104, 13661–13665 (2007).

40. Ieong, K.-W., Uzun, Ü., Selmer, M. & Ehrenberg, M. Two proofreading steps amplify the accuracy of genetic code translation. Proceedings of the National Academy of Sciences 113, 13744–13749 (2016).

41. Goodenbour, J. M. & Pan, T. Diversity of tRNA genes in eukaryotes. Nucleic Acids Res 34, 6137–6146 (2006).

42. Fluitt, A., Pienaar, E. & Viljoen, H. Ribosome Kinetics and aa-tRNA Competition Determine Rate and Fidelity of Peptide Synthesis. Comput Biol Chem 31, 335–346 (2007).

43. Torrent, M., Chalancon, G., de Groot, N. S., Wuster, A. & Madan Babu, M. Cells alter their tRNA abundance to selectively regulate protein synthesis during stress conditions. Science Signaling 11, eaat6409 (2018).

44. Aguilar Rangel, M., Stein, K. & Frydman, J. A machine learning approach uncovers principles and determinants of eukaryotic ribosome pausing. Science Advances 10, eado0738 (2024).

45. Han, Y., Ma, B. & Zhang, K. SPIDER: software for protein identification from sequence tags with de novo sequencing error. J Bioinform Comput Biol 3, 697–716 (2005).

46. Wangen, J. R. & Green, R. Stop codon context influences genome-wide stimulation of termination codon readthrough by aminoglycosides. eLife 9, e52611 (2020).

47. Mangkalaphiban, K. et al. Transcriptome-wide investigation of stop codon readthrough in Saccharomyces cerevisiae. PLOS Genetics 17, e1009538 (2021).

48. Kong, A. T., Leprevost, F. V., Avtonomov, D. M., Mellacheruvu, D. & Nesvizhskii, A. I. MSFragger: ultrafast and comprehensive peptide identification in mass spectrometry–based proteomics. Nat Methods 14, 513–520 (2017).

49. Tsai, A. et al. The Impact of Aminoglycosides on the Dynamics of Translation Elongation. Cell Reports 3, 497–508 (2013).

50. Westhof, E., Yusupov, M. & Yusupova, G. The multiple flavors of GoU pairs in RNA. J Mol Recognit 32, e2782 (2019).

51. Rozov, A., Demeshkina, N., Westhof, E., Yusupov, M. & Yusupova, G. Structural insights into the translational infidelity mechanism. Nat Commun 6, 7251 (2015).

52. Rozov, A., Westhof, E., Yusupov, M. & Yusupova, G. The ribosome prohibits the G•U wobble geometry at the first position of the codon–anticodon helix. Nucleic Acids Res 44, 6434–6441 (2016).

53. Kimsey, I. J., Petzold, K., Sathyamoorthy, B., Stein, Z. W. & Al-Hashimi, H. M. Visualizing transient Watson–Crick-like mispairs in DNA and RNA duplexes. Nature 519, 315–320 (2015).

54. Chen, J. L. et al. Testing the Nearest Neighbor Model for Canonical RNA Base Pairs: Revision of GU Parameters. Biochemistry 51, 3508–3522 (2012).

55. Grosjean, H. & Westhof, E. An integrated, structure- and energy-based view of the genetic code. Nucleic Acids Res 44, 8020–8040 (2016).

56. Behrens, A., Rodschinka, G. & Nedialkova, D. D. High-resolution quantitative profiling of tRNA abundance and modification status in eukaryotes by mim-tRNAseq. Mol Cell 81, 1802–1815.e7 (2021).

57. Tu, L., Khanna, P. & Deutsch, C. Transmembrane segments form tertiary hairpins in the folding vestibule of the ribosome. J Mol Biol 426, 185–198 (2014).

58. Bhushan, S. et al. alpha-Helical nascent polypeptide chains visualized within distinct regions of the ribosomal exit tunnel. Nat Struct Mol Biol 17, 313–317 (2010).

59. Nilsson, O. B. et al. Cotranslational Protein Folding inside the Ribosome Exit Tunnel. Cell Reports 12, 1533–1540 (2015).

60. Freeland, S. J. & Hurst, L. D. The Genetic Code Is One in a Million. J Mol Evol 47, 238– 248 (1998).

61. Jumper, J. et al. Highly accurate protein structure prediction with AlphaFold. Nature 596, 583–589 (2021).

62. Su, J., et al. SaProt: Protein Language Modeling with Structure-aware Vocabulary. in (2023).

63. Hipp, M. S., Kasturi, P. & Hartl, F. U. The proteostasis network and its decline in ageing. Nat Rev Mol Cell Biol 20, 421–435 (2019).

64. López-Otín, C., Blasco, M. A., Partridge, L., Serrano, M. & Kroemer, G. Hallmarks of aging: An expanding universe. Cell 186, 243–278 (2023).

65. Longo, V. D. & Fabrizio, P. Chronological Aging in Saccharomyces cerevisiae. Subcell Biochem 57, 101–121 (2012).

66. Wu, C. C.-C., Zinshteyn, B., Wehner, K. A. & Green, R. High-Resolution Ribosome Profiling Defines Discrete Ribosome Elongation States and Translational Regulation during Cellular Stress. Molecular Cell 73, 959–970.e5 (2019).

67. Cross, K. M. et al. Protein fractional synthesis rates within tissues of high- and low-active mice. PLoS One 15, e0242926 (2020).

68. Pinkard, O., McFarland, S., Sweet, T. & Coller, J. Quantitative tRNA-sequencing uncovers metazoan tissue-specific tRNA regulation. Nat Commun 11, 4104 (2020).

69. Gao, L. et al. Selective gene expression maintains human tRNA anticodon pools during differentiation. Nat Cell Biol 26, 100–112 (2024).

70. Shiraishi, C. et al. RPL3L-containing ribosomes determine translation elongation dynamics required for cardiac function. Nat Commun 14, 2131 (2023).

71. Bernier, C. R., Petrov, A. S., Kovacs, N. A., Penev, P. I. & Williams, L. D. Translation: The Universal Structural Core of Life. Mol Biol Evol 35, 2065–2076 (2018).

72. Zhang, Z., Shah, B. & Bondarenko, P. V. G/U and Certain Wobble Position Mismatches as Possible Main Causes of Amino Acid Misincorporations. Biochemistry 52, 8165–8176 (2013).

73. Franzosa, E. A. & Xia, Y. Structural determinants of protein evolution are context-sensitive at the residue level. Mol Biol Evol 26, 2387–2395 (2009).

74. Drummond, D. A. & Wilke, C. O. The evolutionary consequences of erroneous protein synthesis. Nat Rev Genet 10, 715–724 (2009).

75. Beltran, A., Jiang, X., Shen, Y. & Lehner, B. Site-saturation mutagenesis of 500 human protein domains. Nature 637, 885–894 (2025).

76. Martinez-Miguel, V. E. et al. Increased fidelity of protein synthesis extends lifespan. Cell Metabolism 33, 2288–2300.e12 (2021).

77. Shcherbakov, D. et al. Premature aging in mice with error-prone protein synthesis. Science Advances 8, eabl9051 (2022).

78. Böttger, E. C. et al. Translational error in mice increases with ageing in an organ-dependent manner. Nat Commun 16, 2069 (2025).

79. Di Fraia, D. et al. Altered translation elongation contributes to key hallmarks of aging in the killifish brain. Science 389, eadk3079 (2025).

80. Shahbazian, F. M., Jacobs, M. & Lajtha, A. Rates of protein synthesis in brain and other organs. International Journal of Developmental Neuroscience 5, 39–42 (1987).

81. Stein, K. C., Morales-Polanco, F., van der Lienden, J., Rainbolt, T. K. & Frydman, J. Ageing exacerbates ribosome pausing to disrupt cotranslational proteostasis. Nature 601, 637–642 (2022).

82. Tartaglia, G. G., Pechmann, S., Dobson, C. M. & Vendruscolo, M. Life on the edge: a link between gene expression levels and aggregation rates of human proteins. Trends in Biochemical Sciences 32, 204–206 (2007).

83. Vecchi, G. et al. Proteome-wide observation of the phenomenon of life on the edge of solubility. Proceedings of the National Academy of Sciences 117, 1015–1020 (2020).

84. Hu, J., Wei, M., Mirisola, M. G. & Longo, V. D. Assessing chronological Aging in Saccharomyces cerevisiae. Methods Mol Biol 965, 463–472 (2013).

85. Behrens, A. & Nedialkova, D. D. Experimental and computational workflow for the analysis of tRNA pools from eukaryotic cells by mim-tRNAseq. STAR Protoc 3, 101579 (2022).

86. MGlincy, N. J. & Ingolia, N. T. Transcriptome-wide measurement of translation by ribosome profiling. Methods 126, 112–129 (2017).

87. Langmead, B., Trapnell, C., Pop, M. & Salzberg, S. L. Ultrafast and memory-efficient alignment of short DNA sequences to the human genome. Genome Biol 10, R25 (2009).

88. Lauria, F. et al. riboWaltz: Optimization of ribosome P-site positioning in ribosome profiling data. PLOS Computational Biology 14, e1006169 (2018).

89. Plummer, M. JAGS: A Program for Analysis of Bayesian Graphical Models Using Gibbs Sampling. Proceedings of the 3rd International Workshop on Distributed Statistical Computing (2003).

90. Denwood, M. J. runjags: An R Package Providing Interface Utilities, Model Templates, Parallel Computing Methods and Additional Distributions for MCMC Models in JAGS. Journal of Statistical Software 71, 1–25 (2016).

91. Bertoni, D. et al. AlphaFold Protein Structure Database 2025: a redesigned interface and updated structural coverage. Nucleic Acids Res 54, D358–D362 (2026).

92. Mirdita, M. et al. ColabFold: making protein folding accessible to all. Nat Methods 19, 679–682 (2022).

93. Kabsch, W. & Sander, C. Dictionary of protein secondary structure: Pattern recognition of hydrogen-bonded and geometrical features. Biopolymers 22, 2577–2637 (1983).

94. Su, J. et al. Democratizing protein language model training, sharing and collaboration. Nat Biotechnol 1–7 (2025) doi:10.1038/s41587-025-02859-7.

95. van Kempen, M. et al. Fast and accurate protein structure search with Foldseek. Nat Biotechnol 42, 243–246 (2024).

