## Supplementary material for "Translation fidelity mechanisms and their changes in aging": Methods & supplementary figures

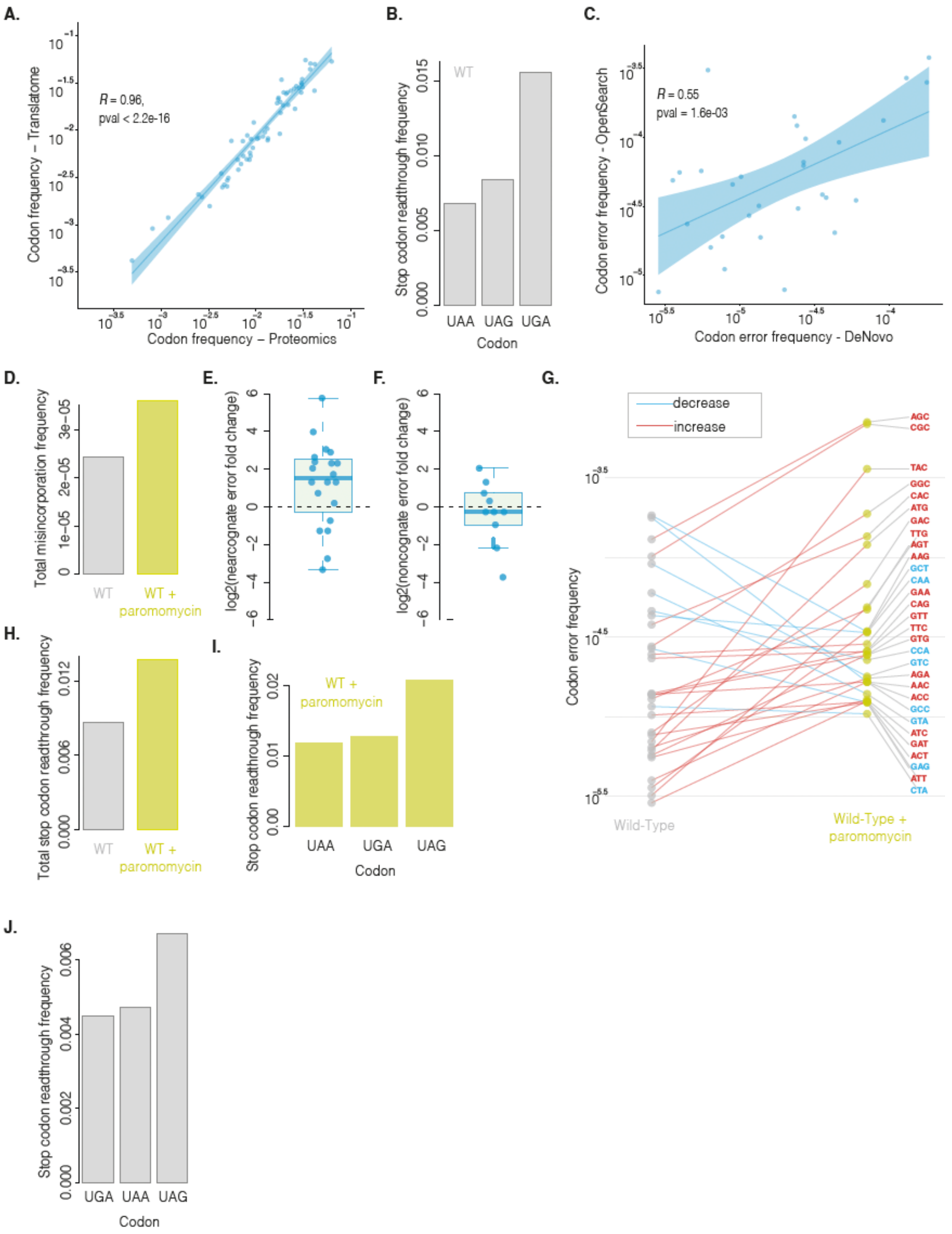


**Figure S1. Technical and biological validation of misincorporation mapping.** A) Correlation plot between codon misincorporation frequencies based on two different spectral assignment strategies: open search with MSFragger, vs. de novo sequencing with PEAKS Online; correlation by Pearson's R. B) Stop codon readthrough frequencies for each stop codon in yeast. C) Comparison between measured codon frequencies in the collected proteomics yeast data and the expected codon composition of the proteome based on ribosome profiling data; correlation by Pearson's R. D) Total misincorporation frequencies for WT yeast and WT yeast cultured with paromomycin. E-F) Near-cognate (E) and non-cognate (F) error frequency fold change in WT+Paromomycin respect to WT. G) Individual error trajectories for codons with measured misincorporation frequencies in both WT and WT+Paromomycin conditions. J) Stop codon readthrough frequencies for each stop codon in mouse.

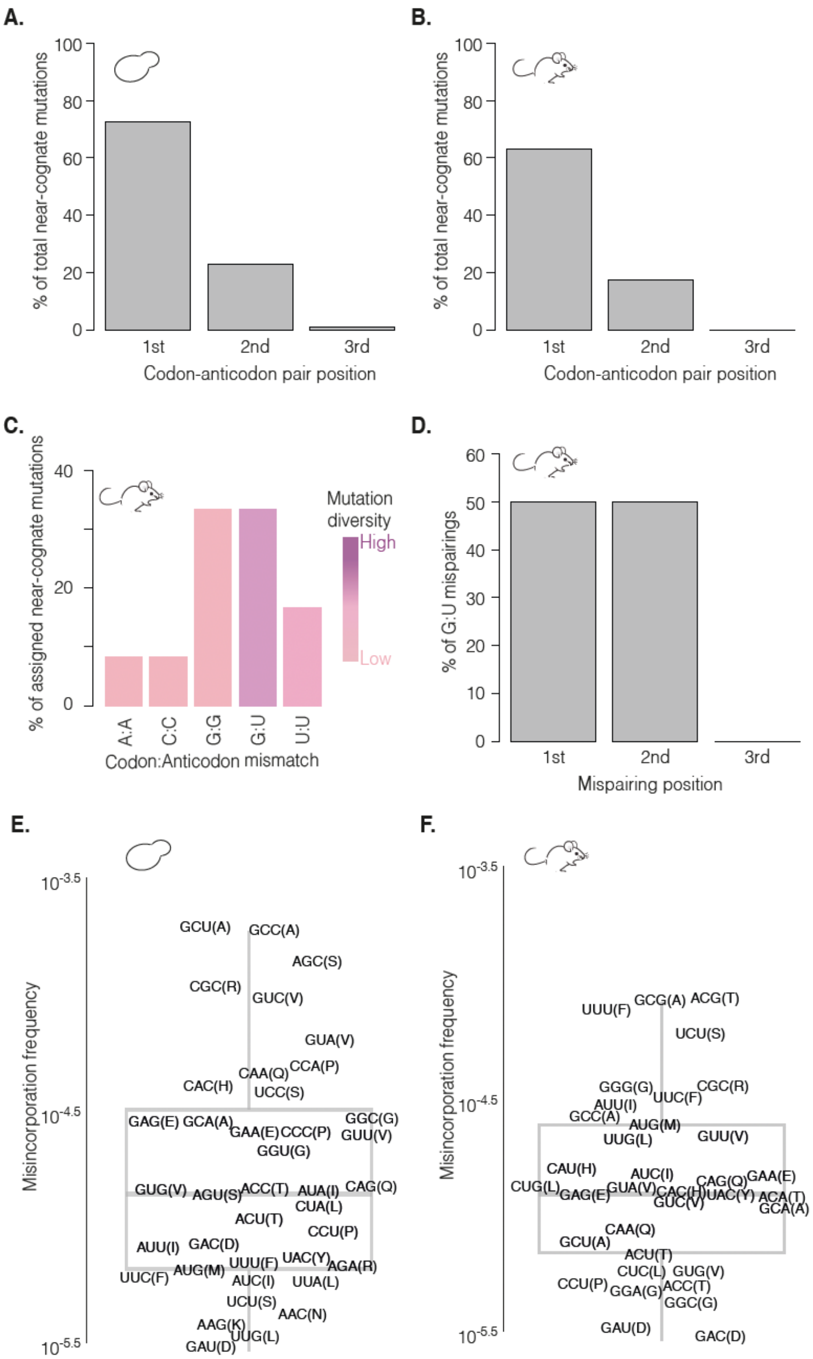


**Figure S2. Features of codon:aticodon mispairings associated to the identified amino acid misincorporations.** A-B) Frequency of misincorporations according to the position of the associated codon:anticodon mispairing in yeast (A) and mouse (B). C) Types and percentages of the different codon:anticodon mispairings assigned to the mapped mouse amino acid misincorporations. D) Percentages of misincorporation-assigned G:U mispairings at each codon:anticodon position, corresponding to mouse misincorporations. E-F) Explicit depiction of codons and their measured error frequencies in yeast (E) and mouse (F).

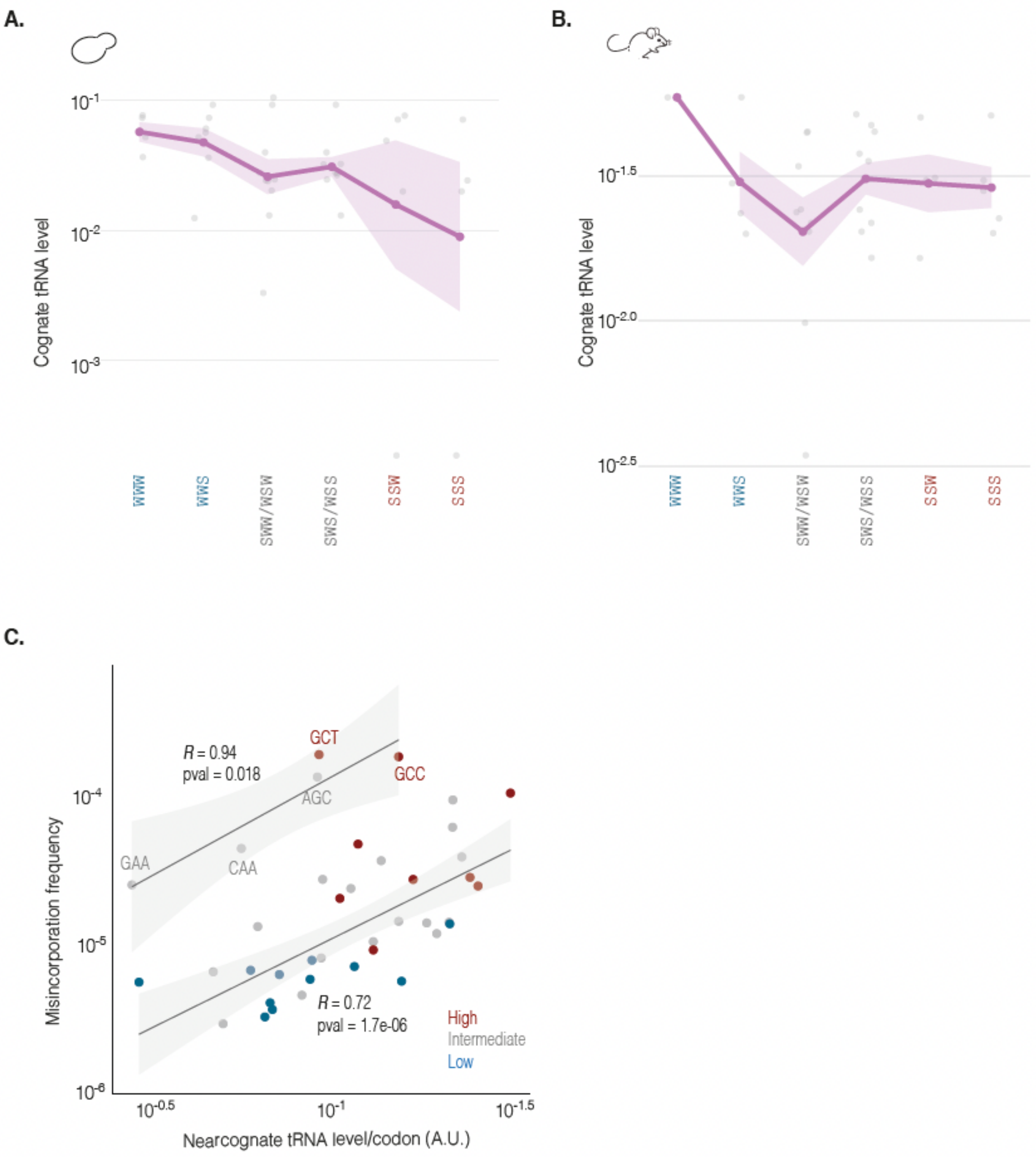


**Figure S3. Cognate tRNA levels according to energy-based codon classification.** A-B) Panels depicting, for the codons with measured misincorporation frequencies in yeast (A) and mouse (B), cognate tRNA levels binned using the energy-based codon classification. Solid line corresponds to the mean and shaded area to the SEM. C) Correlation plot between yeast codon misincorporation frequencies and their corresponding near-cognate tRNA levels over codon usage values. A separate correlation is shown for the group of outliers flagged in Figure 2C. Correlation by Pearson's R.

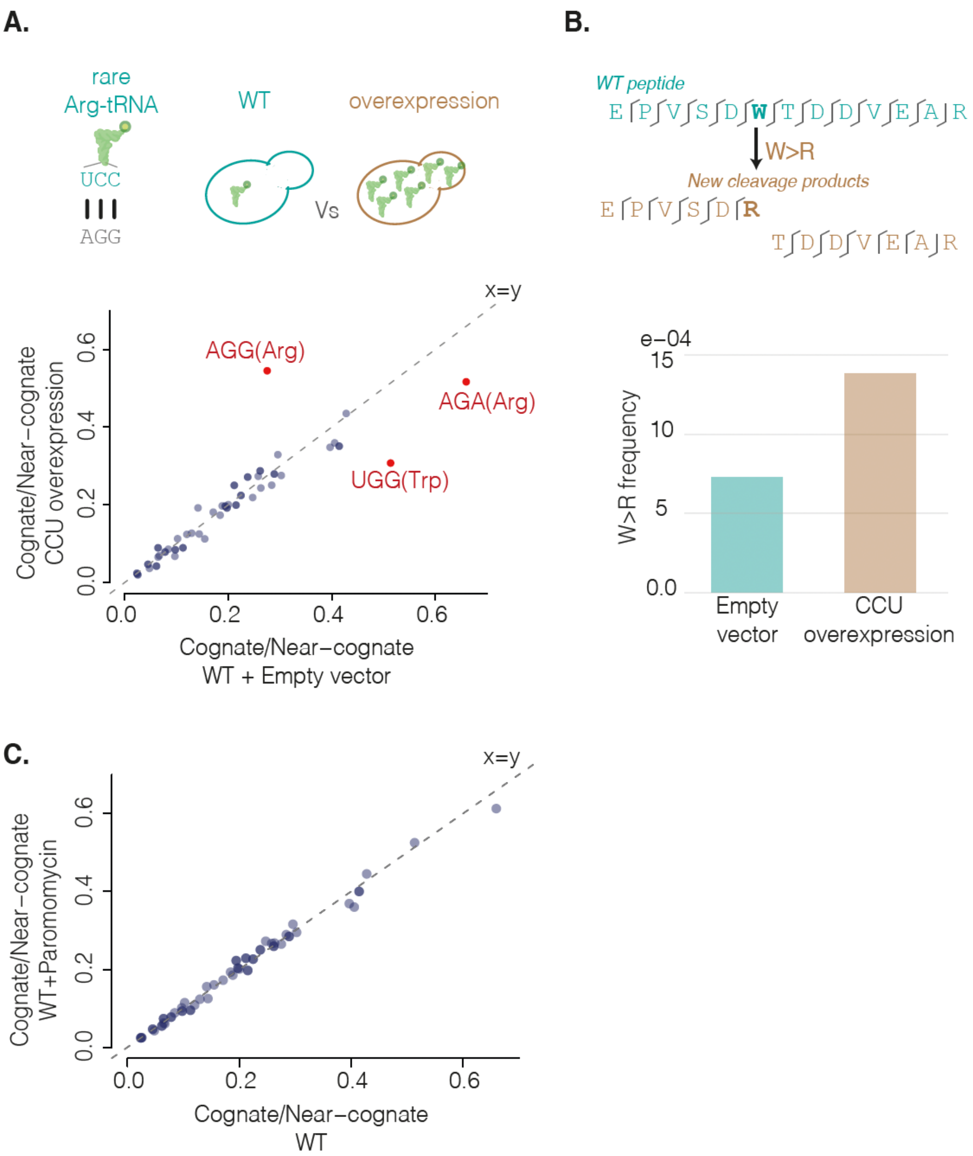


**Figure S4. tRNA pool composition under misincorporation-inducing conditions.** A) Scatter plot comparing cognate:near-cognate tRNA ratios between yeast cells with a plasmid overexpressing $tRNA^{Arg}_{CCU}$ and cells with the empty vector. B) UGG codon misincorporation frequency measured by proxy of W>R mutations in the $tRNA^{Arg}_{CCU}$ overexpression and empty vector yeast strains. C) Scatter plot comparing cognate:near-cognate tRNA ratios between WT yeast cells and WT+Paromomycin.

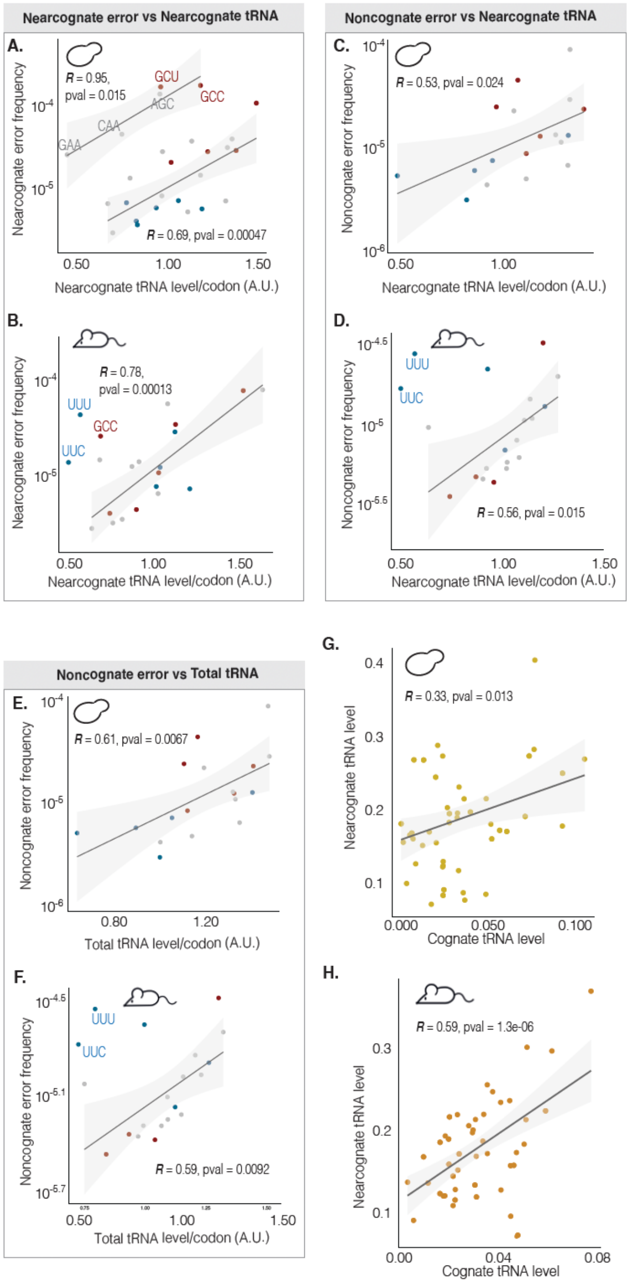


**Figure S5. Non-cognate error frequencies correlate with tRNA pool in yeast and mouse.** A-B) Correlation between codon near-cognate misincorporation frequencies and near-cognate tRNA levels per codon in yeast (A) and mouse (B). C-D) Correlation between codon non-cognate misincorporation frequencies and near-cognate tRNA levels per codon in yeast (C) and mouse (D). E-F) Correlation between codon non-cognate misincorporation frequencies and cognate-plus-near-cognate tRNA levels per codon in yeast (E) and mouse (F). G-H) Correlation between near-cognate and non-cognate tRNA levels in yeast (G) and mouse (H). All correlation by Pearson's R.

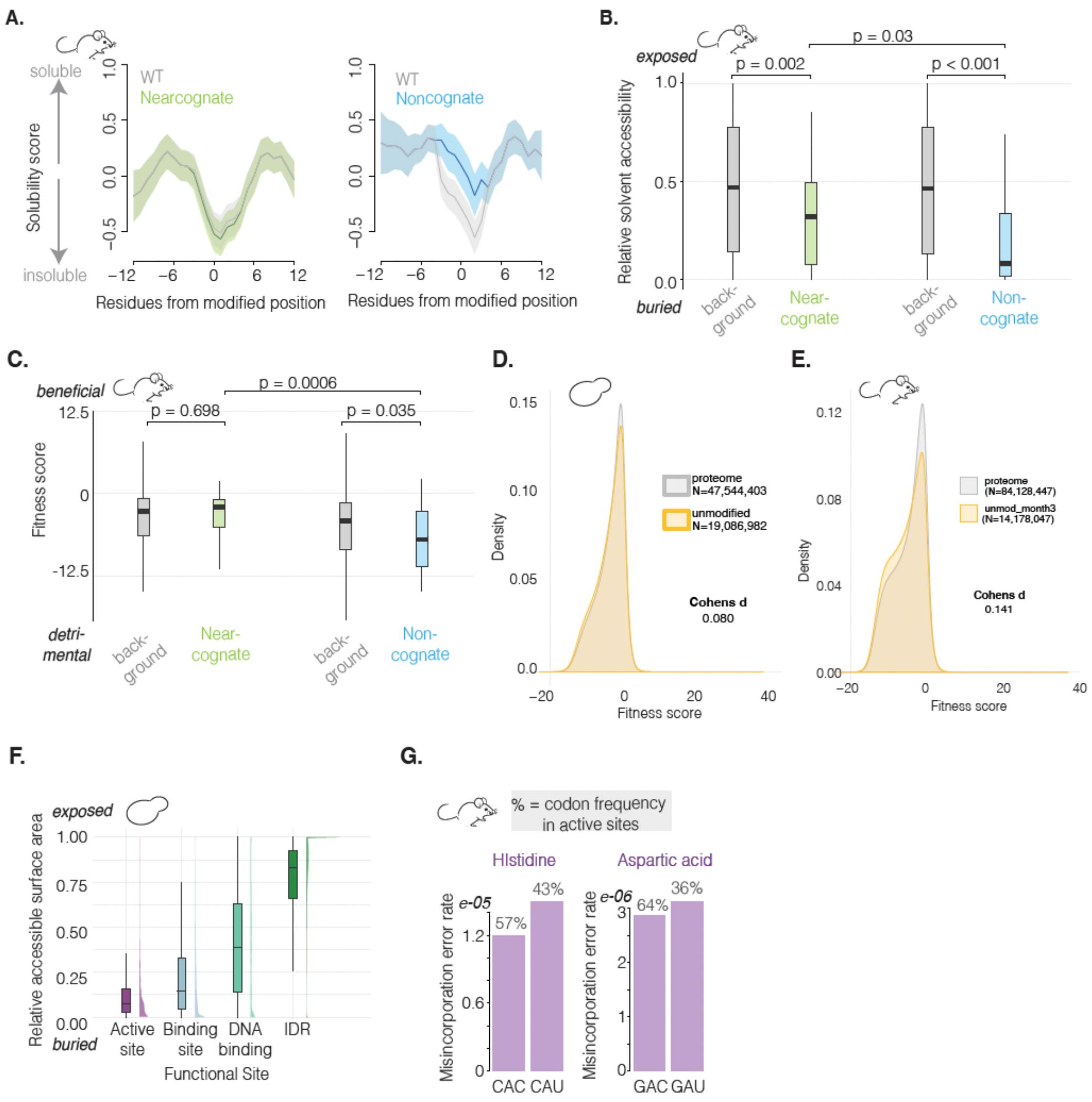


**Figure S6. Structural and functional implications of protein translation errors in mouse.** A) Sequence-based solubility values for all identified misincorporation variants (green – near-cognate, blue non-cognate), and their wild-type counterparts (grey), in their local peptidic context. Misincorporation position centered at position zero, x-axis, flanked by 12 residues at each side; mean (solid line) and SEM (shaded area). B-C) Relative solvent accessibility values (B) and fitness scores (C) for the identified misincorporation variants (green – near-cognate, blue non-cognate), compared to their respective background distributions (i.e. RSAs or fitness scores for all possible near- or non-cognate mutations in the proteome). D-E) Comparison between the distributions of fitness scores for all possible mutations in the measured proteome and the possible mutations sampled by the identified peptides, for yeast (D) and mouse (E). Effect size by Cohen's d. F) Residue solvent accessibility distributions for four different functional classes in the yeast proteome. G) Codon misincorporation frequencies for synonymous codons encoding different catalytic residues. Percentages indicate prevalence at codons encoding catalytic sites.

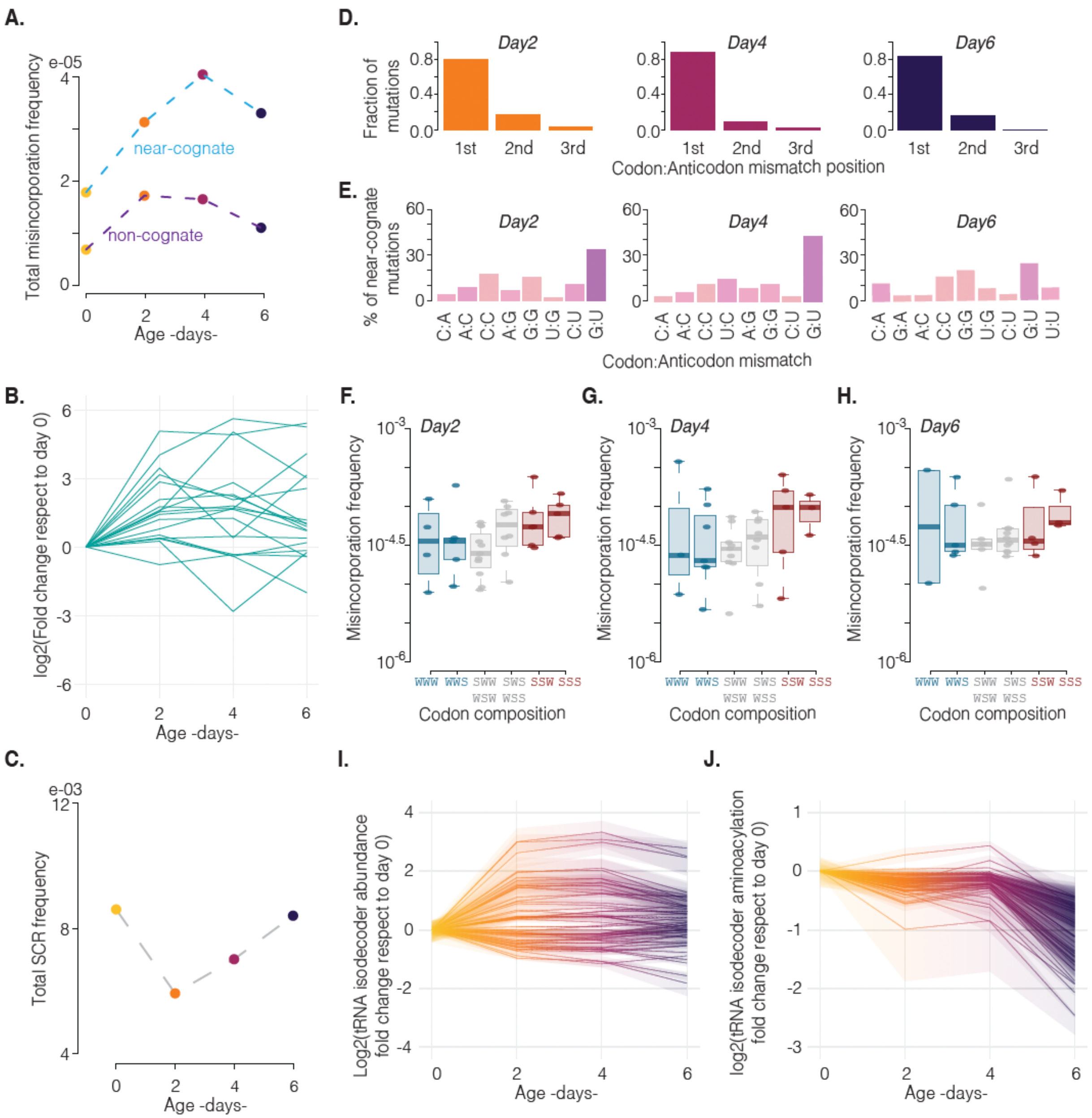


**Figure S7. Protein error trends in aging yeast and their associated codon:anticodon features.** A) Near- and non-cognate amino acid misincorporation frequency as a function of age. B) Fold change, respect to day 0, in codon misincorporation frequencies during yeast aging. C) Total stop codon readthrough frequency as a function of yeast age. D) Frequency of misincorporations according to the position of the associated codon:anticodon mispairing. E) Types and percentages of the different codon:anticodon mispairings assigned to the mapped amino acid misincorporations. F-H) Measured codon misincorporation frequencies binned according to the corresponding energy-based category of the codon. I) Fold change in tRNA isodecor level respect to day 0, each line represents a different isodecoder. J) Fold change in tRNA isodecor aminoacylation level respect to day 0, each line represents a different isodecoder.

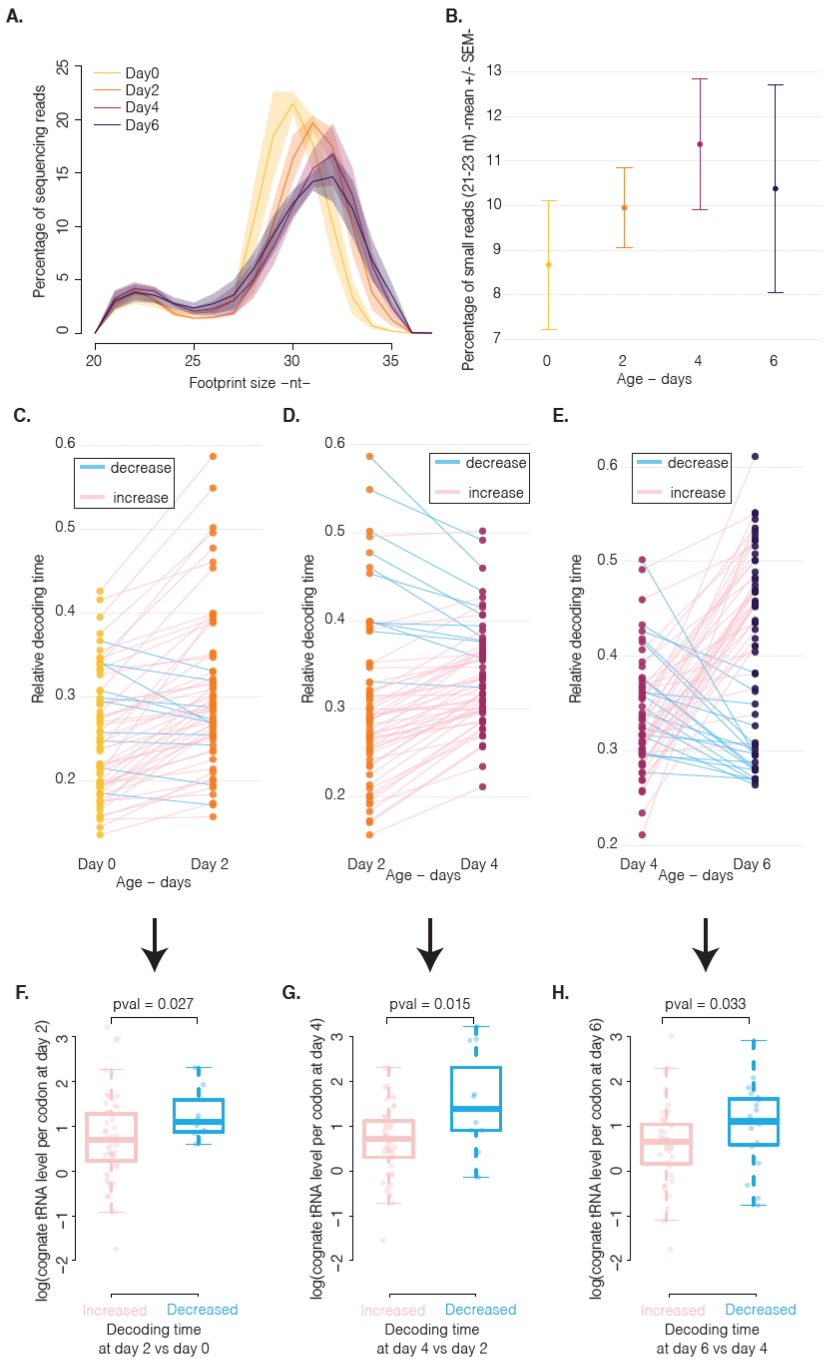


**Figure S8. Codon decoding speed variation as a function of age in yeast.** A) Distribution of ribosome footprint lengths from ribosome profiling libraries of aging yeast samples; mean and SEM. B) Percentage of small ribosome profiling footprints, 21-23nt, in the distributions shown in panel A. C-E) Changes in codon decoding time in consecutive time point pairs, indicating whether decoding time increases of decreases. F-H) Comparison between the aminoacylated cognate tRNA levels per codon for codons whose decoding time increased or decreased as indicated in the corresponding C-E panels.

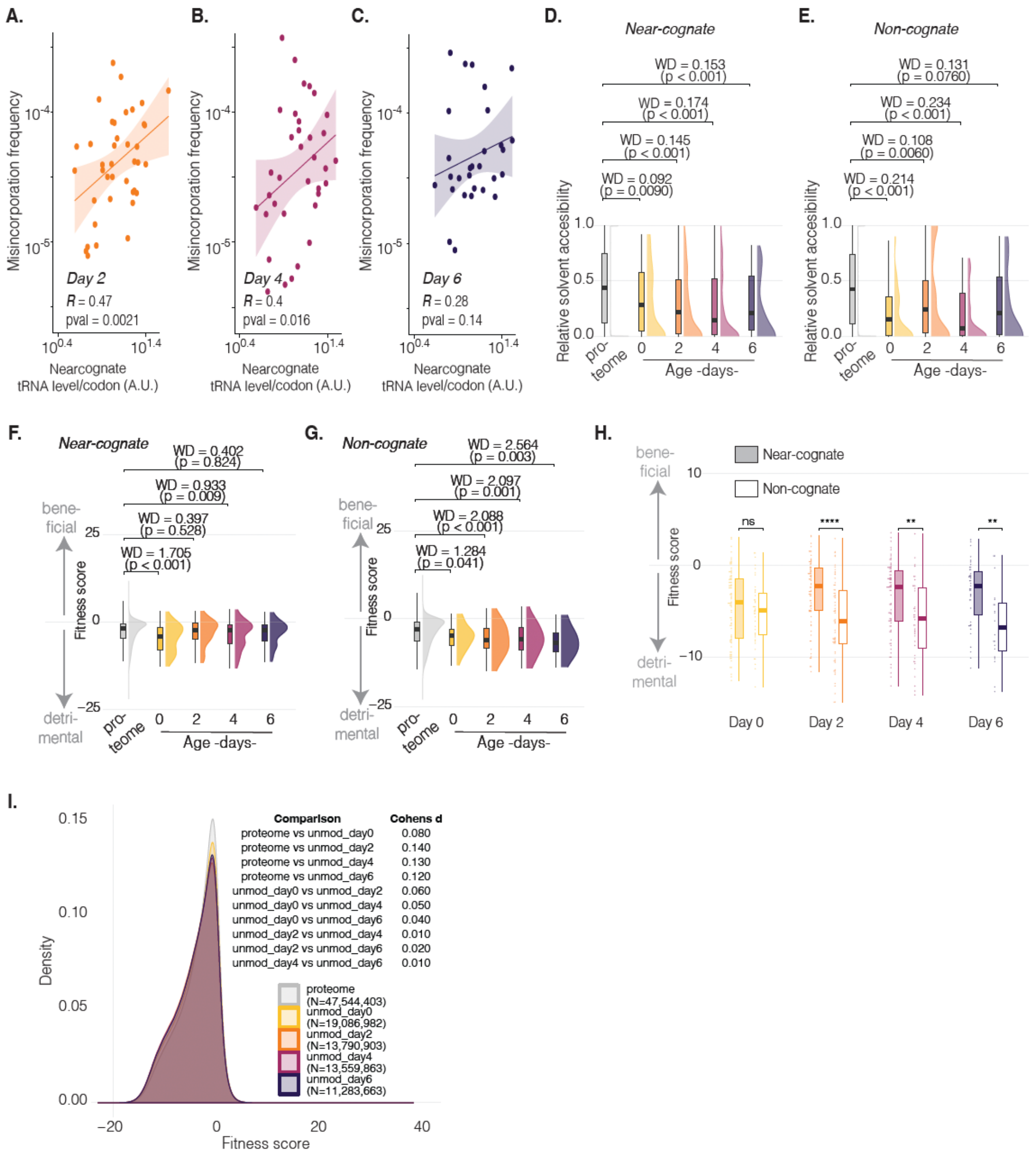


**Figure S9. Misincorporation error frequencies in aging yeast and their structural and functional implications.** A-C) Codon misincorporation frequencies as a function of near-cognate tRNA level per codon, correlation by Pearson's R. D-E) Relative solvent accessibility values for the identified near-cognate (D) and non-cognate (E) misincorporation variants at each time point, compared to the corresponding background distribution (RSA for all possible mutations). F-G) Fitness scores for the identified near-cognate (F) and non-cognate (G) misincorporation variants at each time point, compared to the corresponding background distribution (fitness scores for all possible mutations). H) Paired comparison between fitness scores of near- and non-cognate misincorporation variants at each time point. I) Comparison between the distributions of fitness scores for all possible mutations in the proteome and the possible mutations sampled by the identified peptides at each time point. Effect size by Cohen's d.

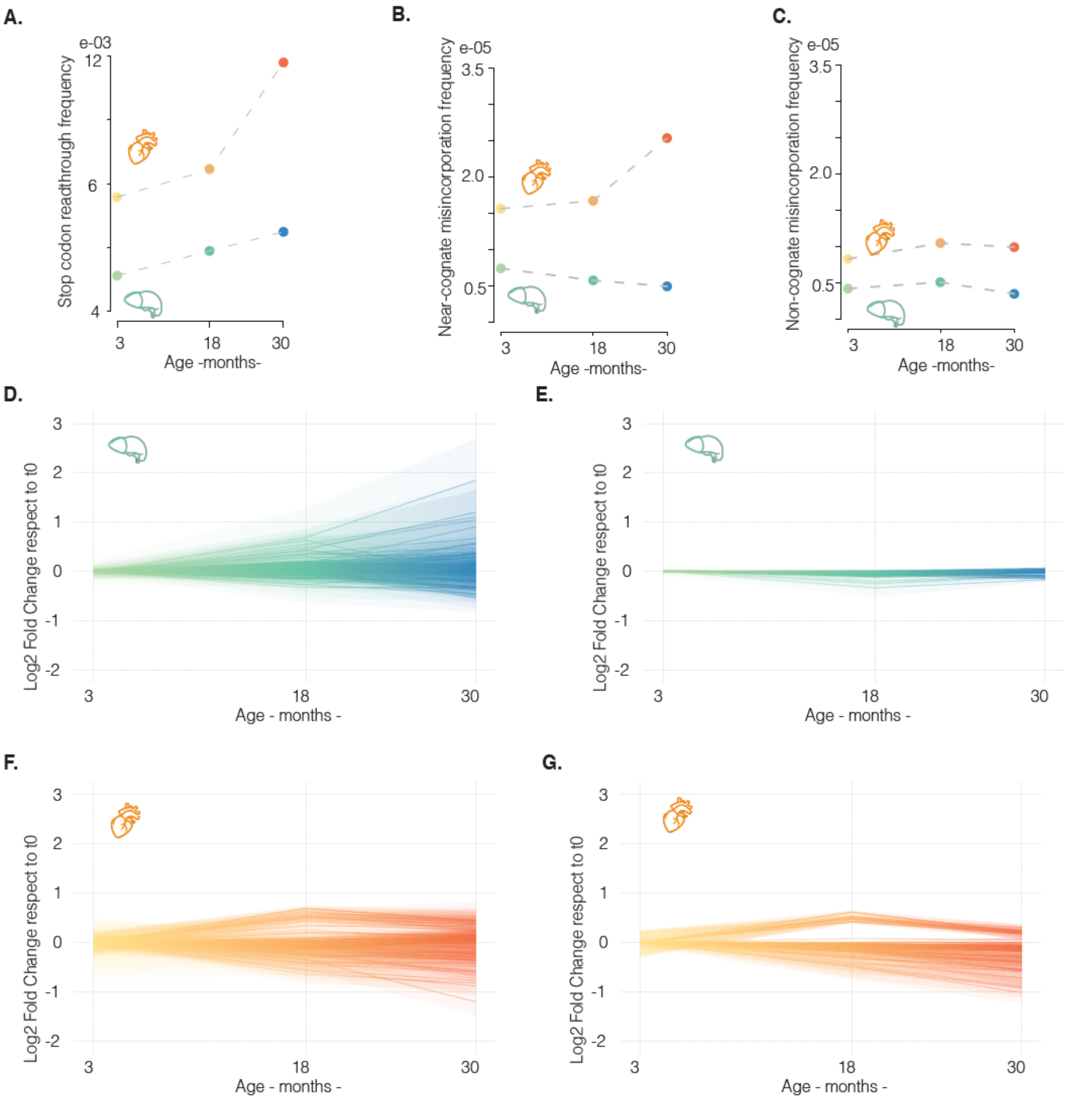


**Figure S10. Organ- and age-dependent variation of protein errors and tRNA pool composition in aging mice.** A) Total stop codon readthrough frequency, as a function of age, for mouse liver (green-blue) and heart (yellow-red). B-C) Near-cognate (B) and non-cognate (C) amino acid misincorporation frequency, as a function of age, for mouse liver and heart. D) Fold change in aminoacylated tRNA isodecoder level in the liver, respect to month 3. E) Fold change in tRNA isodecoder aminoacylation level in the liver, respect to month 3. F) Fold change in aminoacylated tRNA isodecoder level in the heart, respect to month 3. G) Fold change in tRNA isodecoder aminoacylation level in the heart, respect to month 3. In panels D-G each line represents a different isoacceptor.

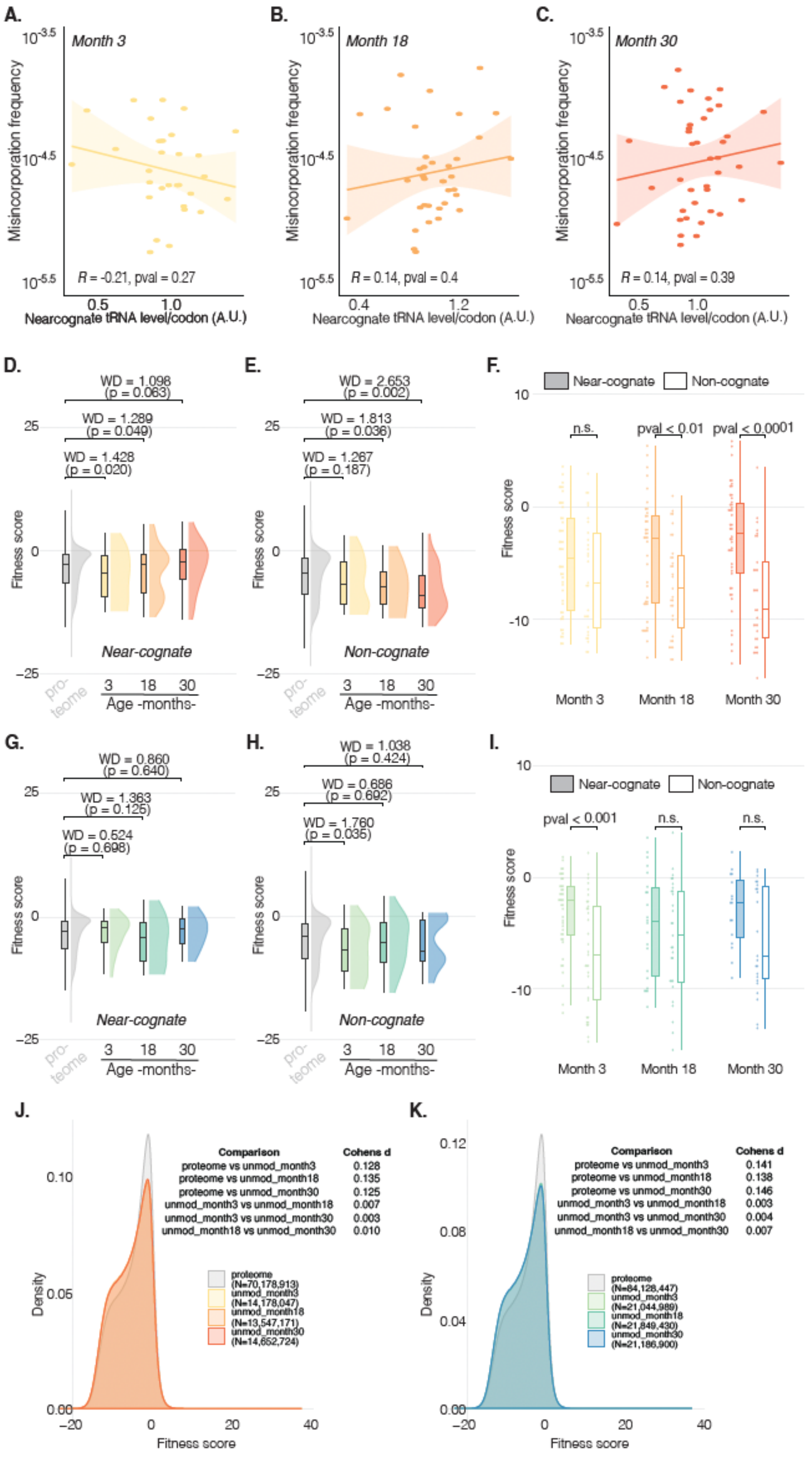


**Figure S11. Misincorporation error frequencies in aging mice and their structural and functional implications.** A-C) Codon misincorporation frequencies as a function of near-cognate tRNA level per codon in hearts of different ages; correlation by Pearson's R. D-E) Fitness scores for the identified near-cognate (D) and non-cognate (E) misincorporation variants in each of the heart samples, compared to the corresponding background distribution (fitness scores for all possible mutations). F) Paired comparison between fitness scores of near- and non-cognate misincorporation variants in each heart sample. G-H) Fitness scores for the identified near-cognate (H) and non-cognate (H) misincorporation variants in each of the liver samples, compared to the corresponding background distribution (fitness scores for all possible mutations). I) Paired comparison between fitness scores of near- and non-cognate misincorporation variants in each heart sample. J-K) Comparison between the distributions of fitness scores for all possible mutations in the proteome and the possible mutations sampled by the identified peptides in each heart (J) or liver (K) sample. Effect size by Cohen's d.

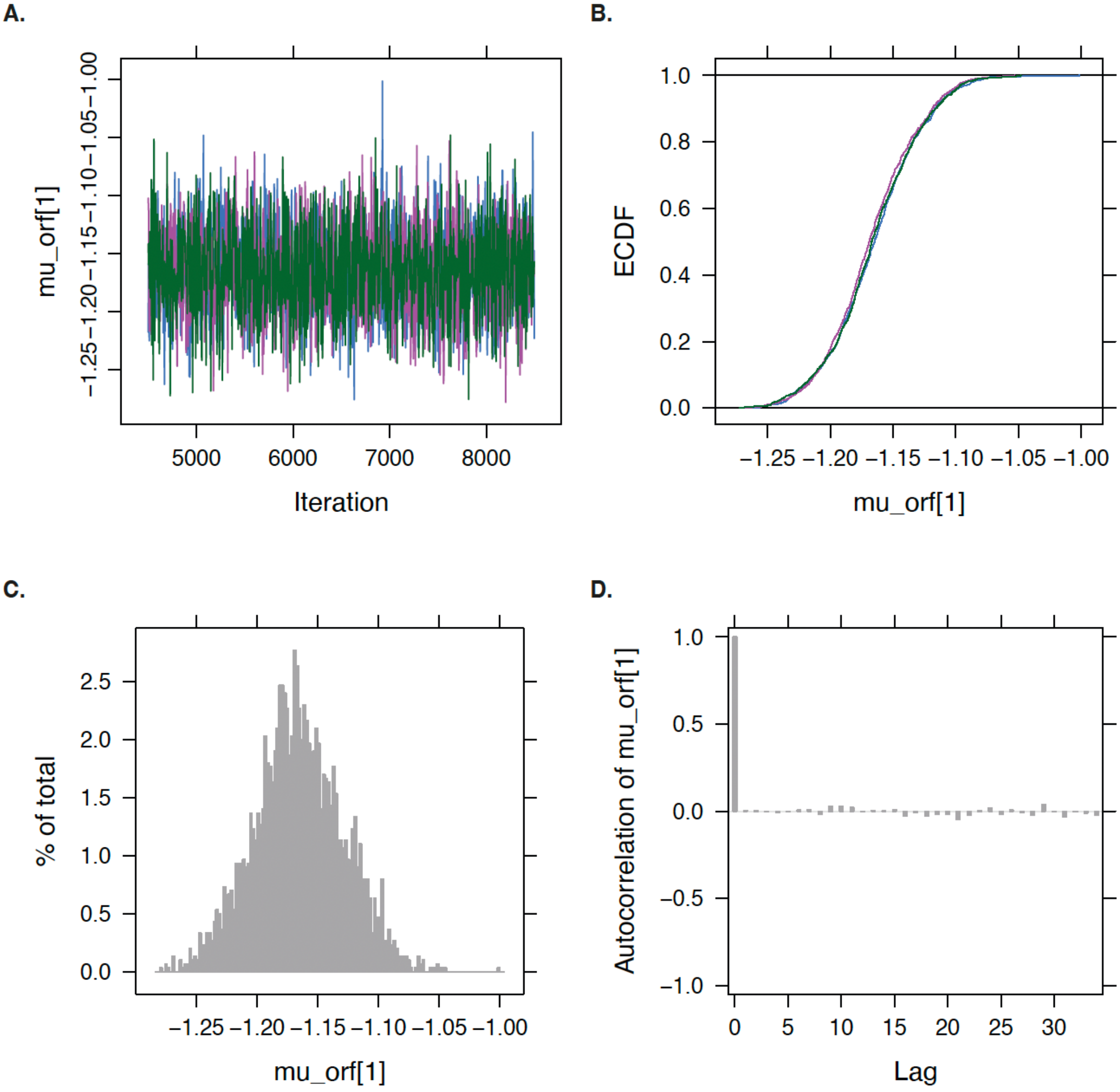


**Figure S12. Example monitoring statistics for ribosome decoding time simulations.** A)Representative trace behavior during chain inference for an individual codon, 3 separate chains – blue, green, magenta. B) Empirical cumulative distribution function overlap of the independent chains. C) Probability density function plot of the estimates. D) Chain autocorrelation plot.

**Table S1. Peptide spectrum match (PSM) breakdown across the data collected in this study.**

| Total PSMs | WT PSMs | PTM-modified PSMs | Candidate substitutions | Accepted substitutions | Near-cognate substitutions | Non-cognate substitutions | Sample |
|---|---|---|---|---|---|---|---|
| **552702** | 518980 | 32571 | 1151 | 204 | 148 | 56 | S.cerevisiae Day 0 |
| **417298** | 381166 | 33797 | 2335 | 263 | 170 | 93 | S.cerevisiae Day 2 |
| **421349** | 379013 | 39979 | 2357 | 329 | 234 | 95 | S.cerevisiae Day 4 |
| **305917** | 275144 | 29261 | 1512 | 177 | 133 | 44 | S.cerevisiae Day 6 |
| **649522** | 621809 | 25945 | 1768 | 358 | 306 | 52 | S.cerevisiae Paromomycin |
| **698783** | 670985 | 26151 | 1647 | 127 | 81 | 46 | M. musculus liver month 3 |
| **697465** | 671506 | 24682 | 1277 | 96 | 51 | 45 | M. musculus liver month 18 |
| **712382** | 688518 | 22783 | 1081 | 75 | 44 | 31 | M. musculus liver month 30 |
| **476854** | 452767 | 22533 | 1554 | 127 | 83 | 44 | M. musculus heart month 3 |
| **455410** | 430645 | 23129 | 1636 | 135 | 83 | 52 | M. musculus heart month 18 |
| **485321** | 458526 | 25070 | 1725 | 192 | 138 | 54 | M. musculus heart month 30 |

**Table S2. Amino acid variant comparison between *S. cerevisiae* s288c and BY4743.**

| Accession | Position | s288c | BY4743 | SNP flagged as misincorporation |
|---|---|---|---|---|
| O13527\|YA11B_YEAST | 138 | L | F | |
| P11154\|PYC1_YEAST | 595 | R | A | |
| P11972\|SST2_YEAST | 644 | E | D | |
| P16862\|PFKA2_YEAST | 880 | K | E | TRUE |
| P24482\|DPB2_YEAST | 458 | F | Y | TRUE |
| P24583\|KPC1_YEAST | 866 | I | L | |
| P32874\|HFA1_YEAST | 1027 | K | E | |
| P33310\|MDL1_YEAST | 150 | F | L | |
| P38781\|RSC30_YEAST | 571 | G | D | |
| P40547\|VID28_YEAST | 758 | N | Y | |
| P53914\|NAT10_YEAST | 856 | G | V | |
| Q00402\|NUM1_YEAST | 895 | S | P | |
| Q03407\|PKH1_YEAST | 187 | F | I | |
| Q03434\|YM12B_YEAST | 629 | R | W | |
| Q04214\|YM13B_YEAST | 360 | H | Y | |
| Q04371\|ARMT1_YEAST | 10 | I | N | TRUE |
| Q12222\|YGK3_YEAST | 59 | I | Y | |
| Q12490\|YB11B_YEAST | 325 | P | R | |
| Q99231\|YD12B_YEAST | 1077 | V | A | |
| P13181\|GAL2_YEAST | 392 | H | R | |
| P33315\|TKT2_YEAST | 632 | F | L | TRUE |
| P39000\|SHC1_YEAST | 233 | K | E | |
| P39106\|MNN1_YEAST | 338 | S | T | |
| P40434\|YIR7_YEAST | 728 | C | R | |
| P53320\|MTM1_YEAST | 343 | R | A | |
| Q03735\|NAB6_YEAST | 77 | F | L | |
| Q06892\|POS5_YEAST | 180 | S | L | |
| Q12751\|RTP1_YEAST | 494 | D | G | |
| P0C2H7\|RL27B_YEAST | 51 | S | L | |
| P47187\|COS5_YEAST | 50 | P | R | |
| Q04215\|YM13A_YEAST | 360 | H | Y | |
| Q06109\|RRG8_YEAST | 251 | N | K | |
| Q12266\|YB11A_YEAST | 325 | P | R | |
| P32389\|MET4_YEAST | 441 | R | A | |
| P32389\|MET4_YEAST | 442 | G | A | |
| P89105\|CTR9_YEAST | 926 | G | E | |
| P89105\|CTR9_YEAST | 927 | W | R | |

### Materials and Methods

Strains and growth conditions.

The *S. cerevisiae* diploid BY4743 (*MATa/α his3Δ1/his3Δ1 leu2Δ0/leu2Δ0 LYS2/lys2Δ0 met15Δ0/MET15 ura3Δ0/ura3Δ0*) strain was used for all yeast experiments described, except the plasmid overexpression of $tRNA^{Arg}_{CCU}$, for which the BY4741 strain was used (*MATa his3Δ1 leu2Δ0 met15Δ0 ura3Δ0*). For chronological lifespan (CLS) experiments, BY4743 was cultured in 2% dextrose synthetic complete media supplemented with a four-fold excess of the amino acids for which the strain is auxotrophic, and cell survival was monitored daily by colony forming units (*82*). Day 0 sample for CLS was collected at an OD600 of 0.8. Paromomycin cultures were grown in similar conditions to Day 0, but medium was supplemented with 0.5 g/l paromomycin. The vector for $tRNA^{Arg}_{CCU}$ overexpression has a URA3 marker, and thus the empty vector and overexpression strains were grown in uracil dropout medium. All cultures were grown at 30°C. For all experiments, samples were harvested by fast filtration, flash frozen immediately in liquid nitrogen.

C57BL/6J mice were shipped from the National Institute on Aging colony. Up to five mice were housed per cage under specific-pathogen-free conditions in the Wu-Tsai Neurosciences Research Institute Veterinary Service Center at Stanford University School of Medicine. The colony room was maintained on a 12:12 h light/dark cycle with food and water provided ad libitum. Mice were euthanized by perfusion with ice-cold phosphate-buffered saline (PBS) following full anesthetization with Avertin (125– 250 mg/kg intraperitoneal injection). All animal care and experimental procedures complied with the Guide for the Care and Use of Laboratory Animals of the National Institutes of Health and were approved by Stanford University's Administrative Panel on Laboratory Animal Care (APLAC).

Mass spectrometry sample preparation and data acquisition.

Yeast and mouse samples were homogenized in lysis buffer (5% sodium dodecyl sulfate (SDS), 50mM Triethylammonium bicarbonate buffer (TEAB) pH 8.5) using 2.8mm ceramic beads (Fisher Scientific) and a bead mill homogenizer. Universal nuclease (Pierce) was added as necessary. Protein concentration was measured by 3-(4-carboxybenzoyl)quinoline-2-carboxaldehyde assay (Thermo Fisher) and 100µg was normalized to 50µL volumes using buffer containing 10% SDS, 100mM TEAB. Samples were subsequently reduced with 10mM DTT, and alkylated with 20mM iodoacetamide, before final acidification with phosphoric acid. Then, samples were loaded onto S-Trap Micro columns (Protifi) according to the manufacturer's protocol, followed by a 2h in-column digestion with Trypsin/Lys-C (Thermo Fisher) at 47°C. The resulting digested peptides were eluted off the S-Trap columns following the manufacturer's instructions. Finally, samples were fractionated using high pH reversed-phase peptide fractionation kit (Pierce), and pairs of non-contiguous samples were pooled, resulting in a total of four fractions per sample.

Samples were analyzed on a timsTOF Ultra (Bruker Daltonics, Germany) coupled to a nanoElute 2 (Bruker Daltonics, Germany) with an average 50 ng sample input. Samples were eluted off a PepSep Ultra XL C18 column (50 cm length x 75 µm ID x 1.5 µm particle size, P/N 1895846; Bruker Daltonics, Germany) at 50 °C connected to a fused silica 10 µm emitter (Bruker Daltonics, Germany) inside a nanoelectrospray Captive Spray source (Bruker Daltonics, Germany) with a 140 min active gradient (2-30% Solvent B in 140-mins, 30-65% Solvent B in 10 mins, 65-95% Solvent B in 0.10 mins, wash at 95% for 14.90 mins; Solvent B: 0.1% formic acid in acetonitrile, Solvent A: 0.1% formic acid). The TIMS 1/K0 mobility range was 0.70-1.30 V•s/$cm^2$, with a ramp time of 180 ms and 100% duty cycle. The source was set to 1500 V, 3.0 l/min dry gas, and 190°C dry temp. DDA windows had a width of 2 m/z to 700 m/z and 3 m/z starting at 800 m/z with a mass range of 300-1700 Da giving an estimated cycle time of 1.18 s using 10 PASEF ramps. The collision energy ramped from 20 to 59 eV at 0.60 to 1.60 V•s/cm$^2$.

tRNA sequencing

To preserve the tRNA aminoacyl moiety, total RNA was recovered under in cold, acidic conditions. Flash-frozen yeast samples were cryogenically milled together with flash-frozen lysis buffer (100mM sodium acetate pH 4.5, 10mM EDTA) using a Mixer Mill (Retsch) for two 1 minute cycles at 30Hz; the resulting powder was thawed right before RNA extraction. Flash-frozen mouse samples were combined with flash-frozen TRIzol reagent (Invitrogen) and homogenized using 2.8mm ceramic beads (Fisher Scientific) and a bead mill homogenizer, in 30s cycles, chilling in ice between cycles. Total RNA, yeast and mouse, was extracted using acidic phenol, pH 4.3 (Sigma Aldrich). Extracted total RNA was ethanol precipitated and resuspended in 50mM sodium acetate pH 4.5, 1mM EDTA. Samples were used to prepare mim-seq tRNA sequencing libraries as described in (*83*), with 3µg of RNA per sample used after periodate oxidation and β-elimination. Reverse transcription was carried out using MarathonRT for 16 hours, reaction buffer 50mM Tris pH 8.3, 200mM KCl, 5mM DTT, 20% glycerol, 2mM MnCl2, 1mM dNTPs, 20U enzyme. Libraries were sequenced on an AVITI instrument (Element biosciences).

Ribosome profiling

Flash-frozen yeast samples were cryogenically milled together with flash-frozen lysis buffer (20 mM Tris-HCl pH 7.5, 140 mM KCl, 1.5 mM $MgCl_2$, 0.5 mM DTT, 0.1mg/ml, cycloheximide, 0.1mg/ml tigecycline, 1% Triton X-100) using a Mixer Mill (Retsch) for 1 minute at 30Hz. Pulverized lysate was thawed in a water bath at room temperature, followed by a 21,000 x *g* centrifugation step at 4ºC for 20 min. Total RNA concentration was measured and 12 A260 units were digested, at room temperature for 25 minutes, with RNAse I (Ambion). Digestion was stopped by adding SUPERase·In (Ambion) and samples were moved to ice. Ribosomes were isolated by centrifugation on a sucrose cushion, 25% w/v, at 72,000 x *g* and 4ºC for 30 minutes. RNA from the ribosomal pellets was isolated by SDS hot-phenol-chloroform RNA extraction. Extracted RNA was ethanol precipitated and used for downstream ribosomal footprint isolation in the ~17-34nt range. Isolated footprints were used for ribosome profiling library preparation as described in (*84*), during which rRNA depletion was carried out using the *S. cerevisiae* ribopools rRNA depletion kit (siTOOLs biotech) according to manufacturer's instructions. Libraries were sequenced on an AVITI instrument (Element biosciences).

Amino acid misincorporation mapping

To identify peptide spectrum matches (PSMs) corresponding to amino acid misincorporations, we used a process-of-elimination strategy. The first part relies on the identification of candidate mutations using PEAKS Online, as described next. First, raw MS/MS spectra were sequenced using, rendering a taxa agnostic space of all candidate PSMs in any data set. Sequenced spectra were searched against the relevant organismal data base (yeast or mouse) plus universal contaminants, using a 1% FDR peptide identification, which reduces the search space by eliminating wild-type (WT) sequences. 20ppm precursor error mass tolerance and 20mDa fragment mass error tolerance were used, with cysteine carbamidomethylation as a fixed modification and methionine oxidation as a variable one. This step produces a set of WT PSMs which will later be used to estimate codon coverages. After this, PSMs were searched for all possible post-translational modifications (PTMs) in the PEAKS built-in data base, allowing for up to 3 variable modifications per PSM. This step produces a set of PTM PSMs which will later be used to estimate codon coverages. Finally, the remaining PSM search space is searched for possible amino acid misincorporations with the SPIDER algorithm, yielding a final list of candidate PSMs harboring a mutation respect to peptides in the organismal database. This list of candidates was further refined as follows, using custom functions which we have made available as an R package (see Data availability). First, PSMs with multiple assigned PTMs or candidate mutations were discarded, except for cysteine carbamidomethylation and methionine oxidation; PSMs matching to contaminants were also removed. Amino acid misincorporation sites not falling at the peptide's termini were requested to be covered by at least one fragment ion pair with a minimal intensity of 2%; if the misincorporation maps to either the N- or C-terminus, there would be only one supporting fragment ion, so the requested minimal ion intensity was increased to >5% for such cases. PSMs with a misincorporation site whose WT counterpart was not detected were discarded, although the R package gives the option to retain them if the user considers them relevant. Since the data has already been directly searched for PTMs, most mass shifts corresponding to such events have been removed from the candidates list. Nevertheless, we carry out a final filtering by comparing the mass shifts of the mutated vs WT peptide and asking whether they could also be explained by shifts annotated in the unimod or uniprot databases, akin to the dangerous modifications approach by Mordret et al. (*6*). In our pipeline, this mainly filters out a few remaining asparagine deamidation events, which we observe are retained as Asn->Asp mutations even in the presence of a large mass shift error. Then, candidate peptides are matched with their corresponding codon sequences, extracted from the translatome. For candidates mapping to more than one protein it is possible to keep them if and only if the codon sequence of those multiple proteins is 100% identical. This is useful for small organisms, like yeast, where the genome duplication gives a series of paralogues with almost identical sequences, and isoform diversity is not big. However, for larger proteomes, like mouse, we recommend running the pipeline with both options and compare the results, since isoform diversity is much more complex. A final filtering step is used to identify overrepresented PSMs, which can bias the calculations for specific codons. First, we compute a background distribution of PSM overrepresentation probabilities based on how frequently a PSM is present times in the WT data. If $X$ represents a unique candidate PSM, and $c$ the total of PSM candidates, we can get assess their relationship by a binomial distribution:

$$X \sim B(c, p)$$

where $p$ comes from the previously calculated background distribution of WT PSMs. The remaining candidate PSMs after filtering are used to obtain codon error counts, i.e. the codon at which the mutation occurs. This will be the numerator of the codon error frequency. To obtain the denominator, i.e. proteomic sequencing coverage, WT PSMs and PTM PSMs are also translated to their codon sequences and used to calculate codon coverages.

Stop codon readthrough (SCR) identification

To identify possible SCR events, first we built a reference SCR database as follows: for each known yeast or mouse transcript, all isoforms included, we extracted the last 40 sense codons, the stop codon, and 120nt downstream of the stop codon; these SCR sequences were translated in the +0 reading frame and translations were truncated after/if a secondary stop codon was found. Then, for each amino acid SCR sequence, the position corresponding to the stop codon was mutated to each of the 20 canonical amino acids, yielding a SCR peptide data base. The SCR peptide data base was concatenated to the corresponding organismal proteomic data base, generating a SCR+Proteome database. This was done since matching PSMs to the WT sequences is necessary to get a reliable peptide FDR estimation. For each mouse or yeast sample, the raw MS/MS data were de novo sequenced using PEAKS Online, followed by database search against the relevant SCR+Proteome database and the universal contaminants database; both steps used a 20ppm precursor error mass tolerance and 20mDa fragment mass error tolerance, with cysteine carbamidomethylation as a fixed modification and methionine oxidation as a variable one. A 1% FDR peptide identification threshold was applied. PSMs with PTMs other than cysteine carbamidomethylation or methionine oxidation were not considered for further analyses. PSMs ending at the C-terminal residue of any WT protein were classified and counted as WT termination events. PSMs mapping to the SCR database and covering the stop codon mutations were counted as SCR events. For each stop codon, its SCR frequency is the quotient of codon specific SCR events/codon specific WT termination events. Total SCR frequency is the quotient of total SCR events/total WT termination events.

<u>Codon usage, cognate tRNA and near-cognate tRNA level estimations</u>

We referred to cognate tRNAs as the any tRNA delivering the correct amino acid by Watson-Crick interactions, allowing for wobble-pairing at the 3rd position. Near cognate tRNAs deliver the wrong amino acid and couple with one mispairing interaction, also allowing for wobble pairing at the 3rd position. Thus, for the *i-th* sense codon, the cognate or near-cognate level ($W_i$) is defined as the sum of the frequencies of the respective cognate or near-cognate tRNA isodecoders $j = 1, \dots, n_i$:

$$W_i = \sum_{j=1}^{n_i} tRNA_{ij}$$

The codon usage for the *i-th* sense codon in the *k-th* transcript is defined as the codon frequency in the transcript, $C_{ik}$, weighted by the transcript abundance in the translatome, in transcripts per million, $TPM_k$. Then, the codon usage for the *i-th* sense codon in the translatome is the sum of the $C_{ik}$ values across the $k = 1, \dots, N$ transcripts in the translatome:

$$CU_i = \sum_{k=1}^{N} TPM_k * C_{ik}$$

<u>Relative codon decoding time estimation</u>

To estimate relative decoding times based on ribosome profiling used the data collected under the presence of CHX and TIG inhibitors. Reads were aligned against *S. cerevisiae* non-coding RNAs using bowtie (*85*); the remaining non-aligning reads were mapped with bowtie to the *S. cerevisiae* translatome, plus/minus 30nt at either transcript end. The resulting translatome mapping reads were A-site and P-site centered using riboWaltz (*86*), with automatic detection of the best extremity for site-offset calculation. The resulting annotated reads encompass short (21-23nt) and long (28-34nt) read populations, where the former represents ribosomes with an empty A-site, pre-accommodation, and the later ribosomes with an occupied A-site, i.e. post-accommodation. Therefore, for any given codon in the translatome, the ratio of short to long footprints represents its relative decoding time. To simultaneously estimate a global relative decoding time for each of the sense codons we used a hierarchical Bayesian model, to allow for the simultaneous inference of codon-specific decoding times while assuming they originate from a common distribution with a global mean (i.e. global decoding time, $\mu_G$). If the variance of the common distribution is small, most decoding times will be similar, whereas a larger variance would indicate a bigger spread amongst codons. For each sense codon $j$, where $j = 1, \dots, 61$, the riboseq experiment gives $i =$

$1, \ldots, n_j$ sequenced positions in the translatome with an associated short to long read ratio, where $Y_{ij}$ denotes the i-th short to long read ratio for the j-th sense codon. Given the nature of the sequencing in the ribosome profiling experiments, we can assume $Y_{ij}$ to be independent and identically distributed (i.i.d), and given their continuous nature, we will assume they follow a normal distribution: $Y_{ij} \sim Normal(\mu_j, \sigma_j)$. Then, sampling for $j = 1, \ldots, 61$ and $i = 1, \ldots, n_j$:

$$Y_{ij} \mid \mu_j, \sigma_j \sim Normal(\mu_j, \sigma_j)$$

where $\mu_j$ comes from the aforementioned common distribution with a global translation decoding mean time $\mu_G$ and its associated standard deviation $\sigma_G$. Thus, the prior for $\mu_j$ will be a two-stage prior, with the first stage being:

$$\mu_j \mid \mu_G, \sigma_G \sim Normal(\mu_G, \sigma_G)$$

where $\mu_G, \sigma_G$ will be described by hyperprior distributions forming the second stage of the $\mu_j$ prior. Given the normal distribution of the global parameters, the hyperpriors can be assigned a normal for the mean, and gamma for the standard deviation reparametrized as precision (1/variance). Thus, hyperpriors in the second stage for $\mu_j$ are:

$$\mu_G \mid \mu_0, \sigma_0 \sim Normal(\mu_0, \sigma_0)$$

$$\frac{1}{\sigma_G^2} \mid a_0, b_0 \sim Gamma(a_0, b_0)$$

where $\mu_0, \sigma_0$ are approximated from the overall data distributions, and $a_0, b_0$ were assigned values of 1 to have a weakly informative prior.

Following a similar rationale, the standard deviation of the decoding time of each codon, $\sigma_j$, will also have a two-stage prior, reparametrizing it first as precision in order to assign it to a gamma distribution as we did for $\sigma_G$. Thus, the first stage for $\sigma_j$ is:

$$\frac{1}{\sigma_j^2} \mid a_G, b_G \sim Gamma(a_G, b_G)$$

where $a_G, b_G$ can be reparametrized in terms of mean and standard deviation as $\mu_{\sigma j}$ and $\sigma_{\sigma j}$:

$$a_G = \frac{\mu_{\sigma j}^2}{\sigma_{\sigma j}^2};\ b_G = \frac{\mu_{\sigma j}}{\sigma_{\sigma j}^2}$$

The $\mu_{\sigma j}$ and $\sigma_{\sigma j}$ hyperparameters can be assigned to weakly informative gamma hyperprior distributions, which we initialized as $\mu_{\sigma j} \sim Gamma(1, 1/3)$ and $\sigma_{\sigma j} \sim Gamma(1, 1/2)$.

The model was implemented in JAGS (*87*) through the R runjags (*88*) interface and solved by MCMC simulations. Each riboseq replicate for each aging timepoint was run separately with three independent chains, each consisting of 1500 iterations during adaptation, 3000 as burn-in, and 4000 for inference. Chain convergence was monitored and assessed by Gelman-Rubin statistic. Fig. S12A-D gives an example of observed convergence. For each codon, the mean of the three biological replicates per time point was used as the final set of codon decoding time estimates.

Ribosome translation speed estimation

For estimation of the translation speed in mouse heart we used public riboseq data from (*68*) retrieved with the gene expression omnibus ID GSE203072, WT samples only. Reads were aligned against *M. musculus* non-coding RNAs using bowtie (*85*); the remaining non-aligning reads were mapped with bowtie to the *M. musculus* translatome, plus/minus 30nt at either transcript end. The resulting translatome mapping reads were A-site centered using riboWaltz (*86*), with automatic detection of the best extremity for site-offset calculation. A-site centered counts were normalized by the transcripts per million (TPM) of the corresponding transcripts, to preclude the calculations from being dominated by highly translated transcripts. Then, for each sense codon $j$, where $j = 1,\ldots,61$, the riboseq data has $i = 1,\ldots,n_j$ sequenced positions in the translatome with an associated normalized counts value, where $Y_{ij}$ denotes the i-th normalized counts value for the j-th sense codon. Next, we calculated codon translation speeds in the same way as the relative decoding time estimations. Since these translation speed values do not come from a short to long read ratio, which intrinsically controls for variation in library depth, after getting the MCMC translation speed estimates for each biological replicate we converted them to z-scores. Then final estimates were taken as the per codon averages from all biological replicates.

Protein structural and fitness analyses

Reference AlphaFold proteomes for S. cerevisiae (UP000002311) and M. musculus (UP000000589) were downloaded from the AlphaFold Protein Structure Database (*59*, *89*). For the BY4743 protein variants described in Table S1, we replaced the reference AlphaFold structures with the BY4743 structures predicted using ColabFold (*90*). Relative solvent accessibilities for each structure were calculated using DSSP through the  biopython wrapper (*91*). Mutation fitess estimation was performed using the zero-shot estimation by the SaProt protein large language model, 650M parameter version (*60*, *92*). The input for prediction was prepared by running the corresponding alphaFold model with Foldseek (*93*), masking low-confidence regions.

Statistical testing

Statistical tests used are indicated in figure legends. For cases where N is small and normality could not be assumed we used Wilcoxon test. For paired comparisons between sets of identified mutations (small N, hundreds) against their proteome-derived background distributions (large N, millions) we used bootstrapped Wasserstein distance to assess sampling bias. For multiple testing, p-values were adjusted using Benjamini-Hochberg procedure. All statistical testing was performed in the R environment.
